# Early diagnosis of Parkinson’s disease: A cross-species biomarker

**DOI:** 10.1101/2021.10.04.462993

**Authors:** David Mallet, Thibault Dufourd, Mélina Decourt, Carole Carcenac, Paola Bossù, Laure Verlin, Pierre Olivier Fernagut, Marianne Benoit-Marand, Gianfranco Spalletta, Emmanuel L Barbier, Sebastien Carnicella, Véronique Sgambato, Florence Fauvelle, Sabrina Boulet

## Abstract

**Background:** Care management of Parkinson’s disease (PD) patients currently remains symptomatic, especially because diagnosis relying on the expression of the cardinal motor symptoms is made too late. Detecting PD earlier therefore represents a key step for developing therapies able to delay or slow down its progression.

**Methods:** We investigated metabolic markers in three different animal models of PD, mimicking different phases of the disease assessed by behavioral and histological evaluation, and in 2 cohorts of *de novo* PD patients (n = 95). Serum and brain tissue samples were analyzed by nuclear magnetic resonance spectroscopy and data submitted to advanced multivariate statistics.

**Results:** Our translational strategy reveals common metabolic dysregulations in serum of the different animal models and PD patients. Some of them were mirrored in the tissue samples, possibly reflecting pathophysiological mechanisms associated with PD development. Interestingly, some metabolic dysregulations appeared before motor symptom emergence, and could represent early biomarkers of PD.

Finally, we built a composite biomarker with a combination of 6 metabolites. This biomarker discriminated animals mimicking PD from controls, even from the first, non-motor signs and very interestingly, also discriminated PD patients from healthy subjects.

**Conclusion:** From our translational study which included three animal models and two PD patient cohorts, we propose a promising composite biomarker exhibiting a high level of predictivity for PD diagnosis in its early phase, before motor symptoms appearance.

**Fundings:** ANR, DOPALCOMP, Institut National de la Santé et de la Recherche Médicale, Grenoble Alpes University.

## Introduction

Parkinson’s disease (PD) is one of the most prevalent neurodegenerative disease in the world, affecting about 1% of adults older than 60 years^1^. From its onset, the pathology evolves progressively and continuously according to three stages i.e. preclinical, prodromal, and clinical, corresponding respectively to the asymptomatic onset of neurodegeneration, non-motor symptoms, and the onset of motor symptoms^2^, finally allowing the diagnosis to be made^3^. Unfortunately, the motor symptoms appear when 70-80 % of the nigrostriatal dopamine system is already lost^1^, precluding any potential neuroprotective intervention^4^. Furthermore, postmortem and histopathological analyses show that the accuracy of current PD diagnosis is only 53% in patients with motor symptoms less than 5 years old^5^. In the early phases of PD, the one defined as “prodromal” is characterized by a panel of non-motor symptoms including neuropsychiatric disorders like apathy, which could schematically defined as a loss of motivation^6^, observed in up to 70% of PD patients^7^. Although these symptoms are often premonitory of PD, they are not specific and therefore cannot be considered as predictive criteria. Thus, finding reliable, specific and highly predictive biomarkers of the prodromal stage of the disease is a substantial challenge for the success of PD curative or preventive strategies, and for the development of new therapies able to delay or slow down PD progression^8, 9^.

PD is a multifactorial disease^10^ resulting from a complex interplay between genetic and environmental factors. The metabolome - the global pool of metabolites - reflects the interactions between genotype, lifestyle, diet, drug therapy or environmental exposure^11^. Therefore, metabolomics, i.e. metabolome analysis, could represent a powerful tool to elucidate the molecular mechanisms involved in PD and to identify potential predictive biomarkers^12–15^. Metabolic alterations have already been described in patients expressing neuropsychiatric symptoms similar to those observed in the prodromal stage of PD^16^. Additionally, it has been shown that the metabolic dysregulation observed in serum can accurately discriminate newly diagnosed PD patients from controls^17^. Similarly, alterations in plasma metabolome have been correlated with disease progression in PD patients^18^. However, the use of metabolomics as a predictive tool during the prodromal stage remains to be investigated.

To study this prodromal stage, animal models of PD have been largely developed^19^, but none is currently able to recapitulate all phenotypic and etiological characteristics of the disease. We therefore investigated potential metabolic changes in three different animal models, two complementary rodent models and one non-human primate model, expressing complementary PD characteristics^20–22^. We first used the 6-hydroxydopamine (6-OHDA) model as it represents a gold standard and phenotypically correct model of PD that can stably mimic the different stages of the disease including the prodromal phase, without temporal evolution^19, 20^. Moreover it presents a good predictive value regarding the treatments classically used in clinics, such as dopaminergic agonists^23^. We then used a viral vector-induced alpha-synuclein rat model. which presents synucleinopathy and offers the advantage of longitudinal studies thanks to the progressive expression of the vector. Nonetheless this model does not encompass the whole neuropsychiatric symptoms characteristic of prodromal stage of PD (e.g. apathy)^24^. Finally, we used non-human 1-methyl-4-phenyl-1,2,3,6-tetrahydropyridine (MPTP) primates which share greater genome sequence identity with humans and significant neuroanatomical similarities (Figure 1). Metabolic changes were analyzed in blood (serum) to find biomarkers that are easily transposable to the clinic, and in specific brain regions differently affected by the neurodegenerative process (i.e. the dorsal striatum (DS) and nucleus accumbens (Nacc)) to potentially associate these biomarkers to the pathophysiological processes involved in PD. We thereby built a blood biomarker from a combination of metabolites that might be relevant to the prodromal stage. From a translational perspective, and in order to validate the clinical relevance of our PD biomarker candidate, we confronted our pre-clinical results with the metabolomic results of *de novo* PD patients from two different biobanks.

**Figure 1:**
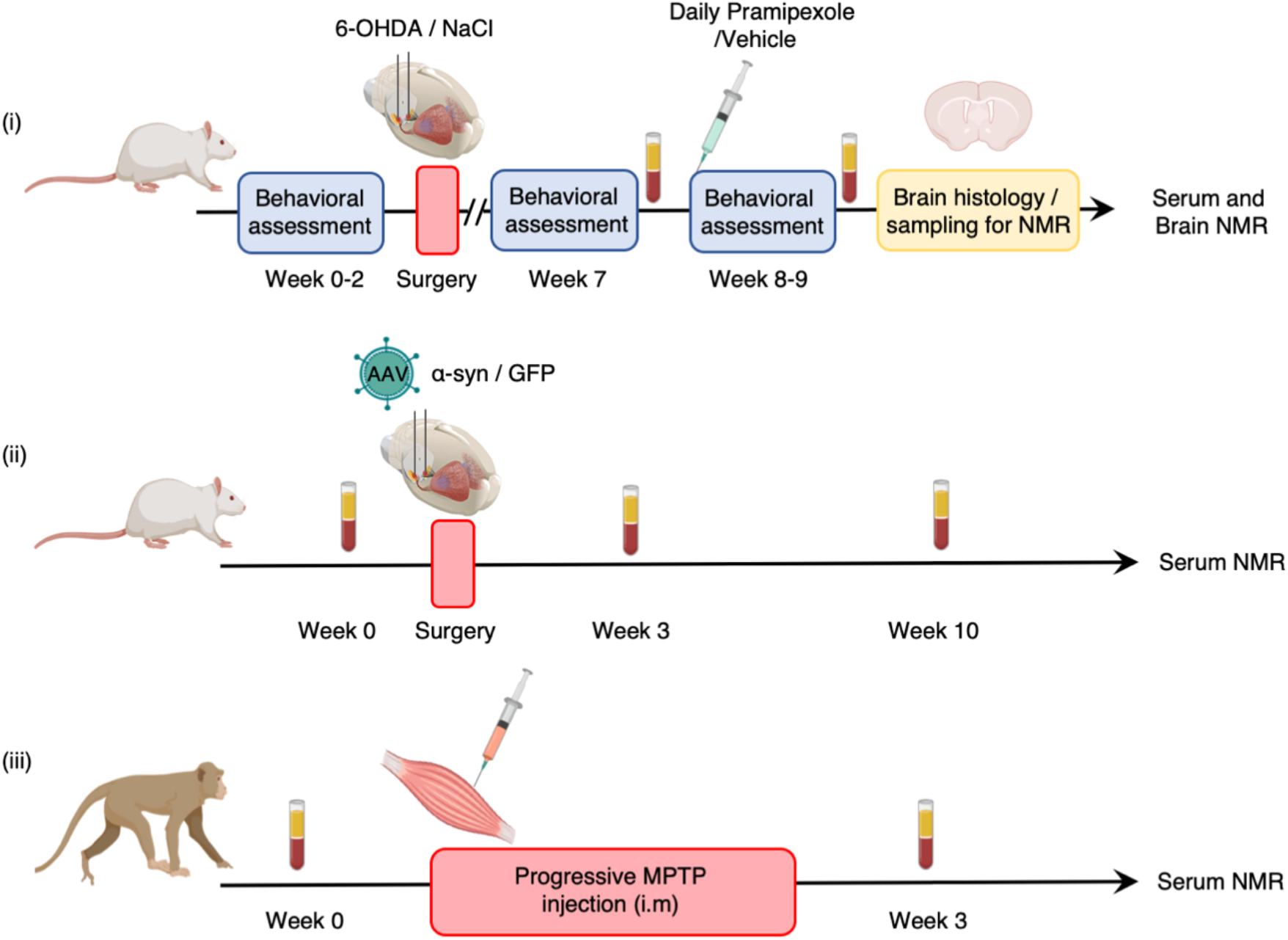
Flowchart of experimental procedures applied to different PD animal models. All serum samples were collected after fasting and following the same protocol. They were analyzed by ^1^H-NMR at 950 MHz. Brain samples of 6-OHDA rats were used for histology and metabolomics analysis performed by ^1^H HRMAS NMR at 500 MHz.

## Results

### The 6-OHDA rat model allows the study of different stages of PD

Although PD diagnosis in human relies on a scale which integrates evaluations of both neuropsychiatric and motor symptoms^25^, its final confirmation is based only on post-mortem histological evaluations (mainly of neuronal loss in the nigro-striatal pathway leading to denervation of DS). In order to characterize the clinical-like progression of the disease in 6-OHDA rats, we established a score, called the Parkinson Disease Progression (PDP) score, based on the same kind of criteria, i.e. (i) the neuropsychiatric component, evaluated by operant self-administration performances (motivation); (ii) fine motor capacities, evaluated by stepping test performances; (iii) the extent of the DS lesion, evaluated by a postmortem histological analysis. Based on this score, we assigned the PD animals to two categories, namely prodromal-like or clinical-like. These terms will be used throughout the study (Figure 2A).

**Figure 2:**
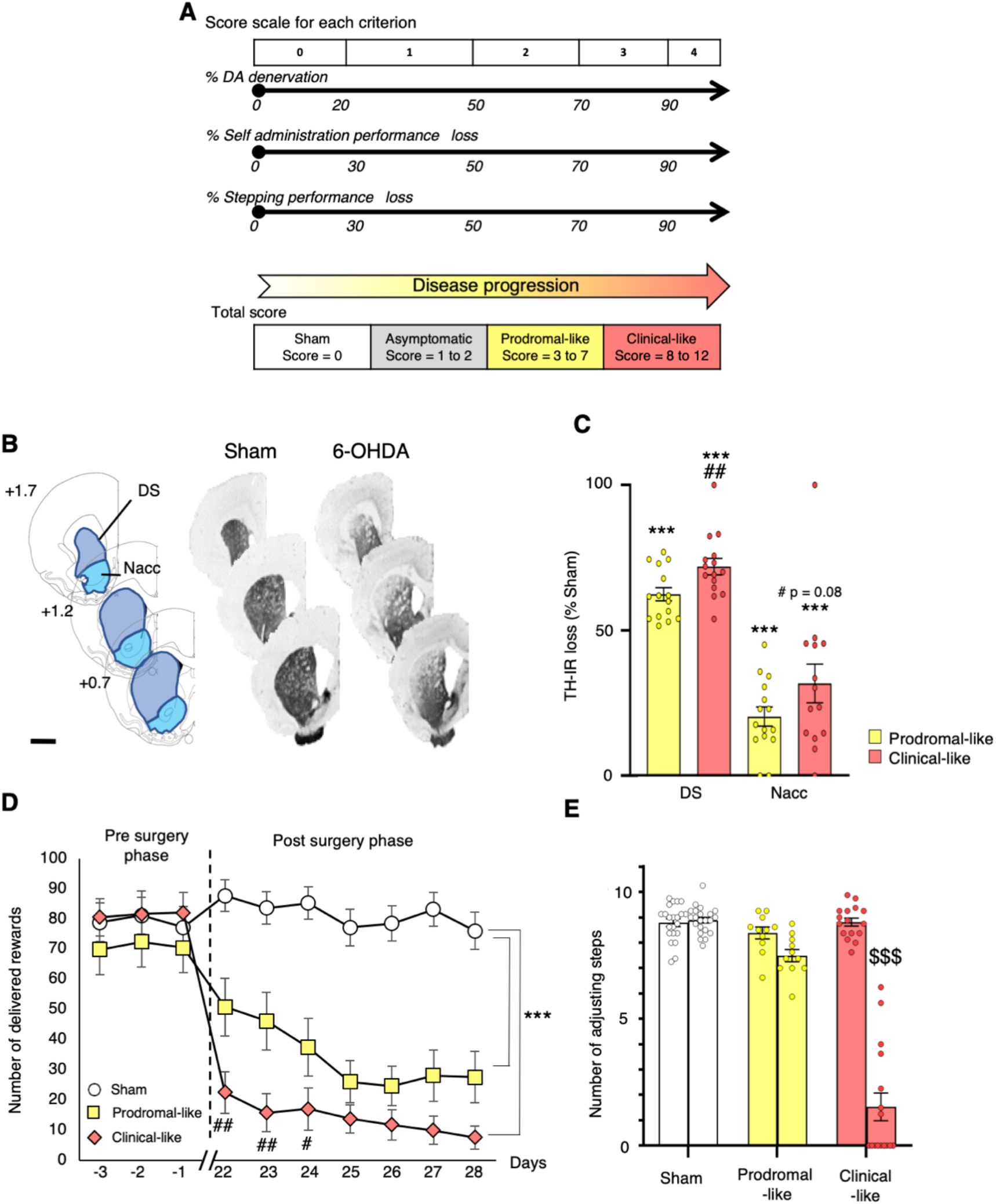
Striatal dopaminergic denervation induced by bilateral 6-OHDA lesion of the SNc leads to apathetic like behavior and fine motor dysfunctions. (A) PDP score used to classify 6-OHDA animals for the disease progression. PDP score is based on the sum of histological and behavioral components (self-administration and stepping performances), taking values from 0 to 4 for each. In this way, a score could be assigned to each animal from 0 (sham animal) to either 3-7 (prodromal-like animal), or 8-12 (clinical-like animal). (B) Examples of coronal sections of sham and 6-OHDA rat brains stained for TH at three representative striatal levels with respect to bregma, and corresponding diagrams selected from the atlas of Paxinos and Watson^79^. Areas used for the quantification of dopaminergic denervation in the different subregions analyzed are illustrated. Scale bar represents 2mm. (C) Quantification of TH-IR staining loss at the striatal levels shown in (B), expressed as a percentage of the mean value obtained for sham-operated animals (n = 22). We observed a large decrease of TH-positive neurons in DS and a lighter decrease in Nacc of prodromal (n = 14) and clinical-like (n = 15) rats. (D) 6-OHDA SNc lesion induced an abrupt instrumental deficit in an operant sucrose self-administration procedure. Results are expressed as the mean number of sucrose deliveries per session. (E) 6-OHDA SNc lesion reduced the number of adjusting steps in a stepping procedure only in clinical-like animals. Results are expressed as the mean number of forelimb adjustments for two trials before (left bar) and after (right bar) 6-OHDA (prodromal- like and clinical-like animals) or saline (sham) injection. Data are presented as means ± SEM and tested by 1-way ANOVA or RM-ANOVA followed by post-hoc test of Tukey or Sidak. ***: p ≤ 0.001 compared to sham; ##: p ≤ 0.01 clinical-like compared to prodromal-like; $$$: p ≤ 0.001 before surgery compared to after surgery.

Tyrosine hydroxylase immunoreactivity (TH-IR) revealed that bilateral 6-OHDA injection in the substantia nigra pars compacta (SNc) led to a partial nigrostriatal dopaminergic lesion resulting in dopaminergic denervation in the DS and, to a lesser extent, in the Nacc (Figure 2B and 2C). Indeed, TH-IR quantification showed a significant loss (62.2%) of dopaminergic projections in the DS of prodromal-like animals compared to shams, with a greater loss in clinical-like animals (71.96%). A slight loss of TH-IR was also observed in the Nacc of 6-OHDA animals (Figure 2C). This denervation pattern preserves learning and global ambulatory activity of animals^20^, allowing study of motivational processes without the potential bias related to locomotor alterations often present in PD animal models (Figure S1)^20, 23^. Motivation was measured by the sucrose self-administration procedure. Prior to surgery, rats learned the motivation task and reached their maximum performance level (about 80 rewards per one-hour session). During the post-surgery phase (i.e. after surgery), the performances of sham rats remained stable, while that of 6-OHDA animals dramatically decreased (30 to 66%) (Figure 2D).

Regarding fine motor skills, the 6-OHDA infusion did not reduce the number of adjusting steps, except in the clinical-like group where it strongly decreased (Figure 2E).

We then investigated the metabolic dysregulations of each animal in association with its PDP score, i.e. its PD stage.

### Serum metabolic signatures co-evolve with PD progression in 6-OHDA rats

Figure 3A illustrates a typical proton nuclear magnetic resonance (^1^H NMR) spectrum of 6-OHDA rat serum, acquired at ultra-high magnetic field (23T). This allowed the identification of approximately 50 metabolites (Table S2).

**Figure 3:**
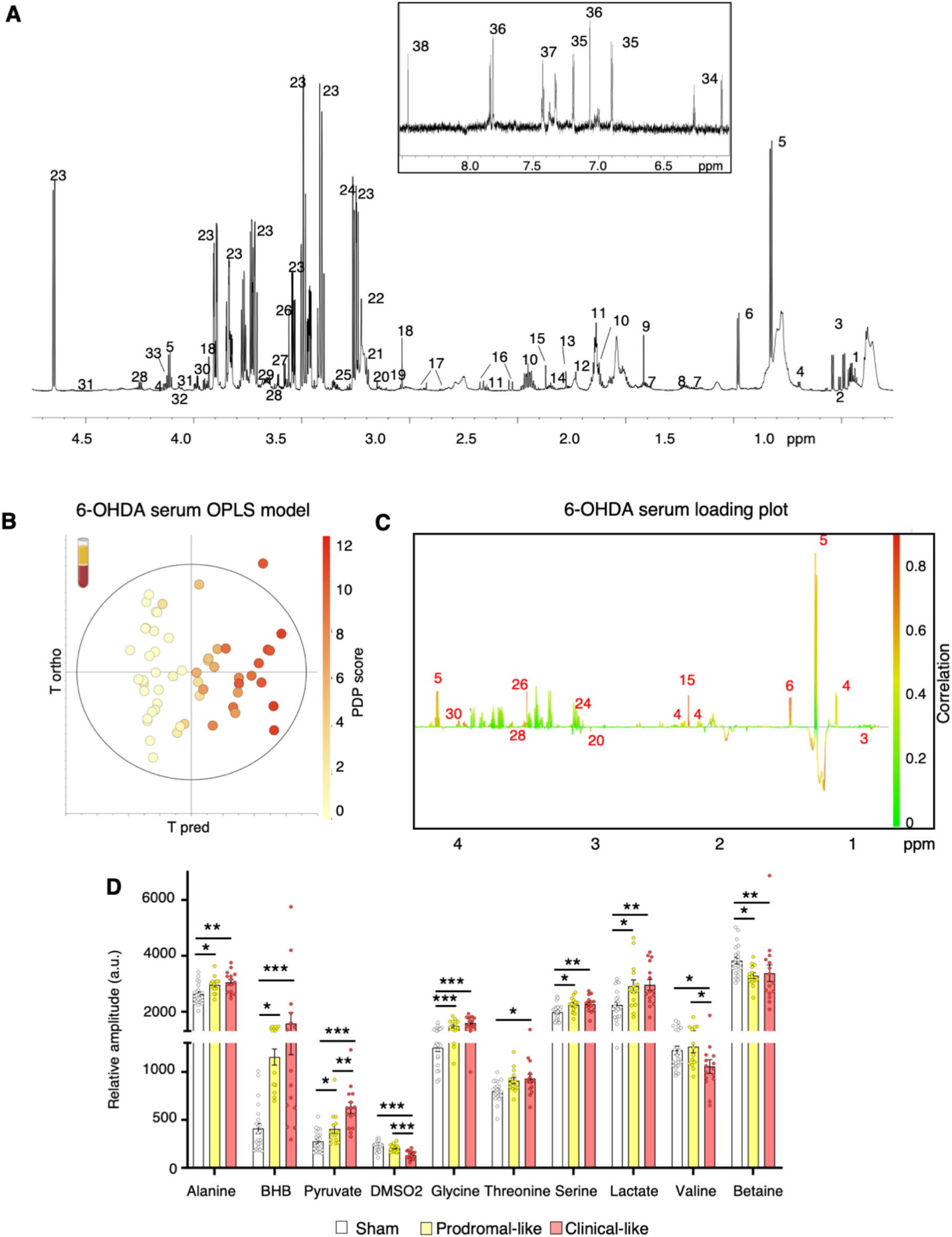
Serum metabolic profile of 6-OHDA rats is gradually modified by PD progression. (A) Example of ^1^HNMR spectrum at 950 MHz (δ0.5–4.7 ppm and δ6–8.5 ppm) using CPMG pulse sequence. Assignment: 1-isoleucine; 2-leucine; 3-valine; 4-BHB; 5-lactate; 6-alanine; 7-arginine; 8-lysine; 9-acetate; 10-glutamine; 11-methionine; 12-acetone; 13-acetoacetate; 14-glutamate; 15-pyruvate; 16-citrate; 17-asparagine; 18-creatine; 19-phosphocreatine; 20-DMSO2; 21-choline ; 22-phosphocholine; 23-glucose; 24-betaine; 25-myo-inositol; 26-glycine; 27-glycerol; 28-threonine; 29-glycerophosphocholine; 30-serine; 31-ascorbate; 32-glycerate; 33-proline; 34-deoxycytidine triphosphate; 35-tyrosine; 36-histidine; 37-phenylalanine; 38-formate. Macromolecules are not specified (see supplemental Table2). (B-C) OPLS model built with ^1^HNMR spectra of serum samples from 6-OHDA (n = 29) and sham (n = 22) rats, and their PD scores: the “6-OHDA serum OPLS model”. R2Y = 0.926; Q^2^ = 0.604; 1 predictive and 3 orthogonal components; CV-ANOVA *p-*value = 3.47 x10^-7^. (B) Score plot *vs* the first predictive and first orthogonal components. A clear gradation of color is observed from the left to the right, showing that metabolic profiles evolve with PD progression. (C)Loadings plotted in 1 dimension with NMR variables color coded for their correlation with PD score from green (low correlation) to red (high correlation). Positive peaks indicate up-regulated metabolites along with increasing PDP score, while negative peaks indicate down-regulated metabolites along with PDP score evolution. (D) Relative amplitude of metabolites the most involved in metabolic gradation observed in 6-OHDA animals, i.e. alanine, betaine, BHB, DMSO2, glycine, lactate, pyruvate, serine, threonine, valine, in sham (n = 22), prodromal-like (n=14) and clinical-like (n = 15) animals. Mean ± SEM, one-way ANOVA followed by post-hoc test of Tukey and correction for multiple comparisons. *: p ≤ 0.05; **: p ≤ 0.01; ***: p ≤ 0.001 *a.u: arbitrary units*

The score plot of the orthogonal partial least squares (OPLS) analysis, performed with the NMR data and PDP scores for each animal (i.e. the “6-OHDA serum OPLS model”) shows a clear gradation of dot colors from “cold” (white) on the left to “hot” (red) on the right. This indicates that the metabolic profile co-evolves with disease progression (Figure 3B). Among the 14 most relevant metabolites (i.e. correlation ≥ 0.5), ten were significantly modified in at least one PD-like group compared to sham animals (Figure 3C): alanine, betaine, β-hydroxybutyrate (BHB), dimethyl sulfone (DMSO2), glycine, lactate, pyruvate, threonine, serine, and valine. These metabolites did not evolve identically with disease progression. First, 6 metabolites were significantly modified in the prodromal-like stage: alanine, BHB, glycine, lactate and serine increased, while betaine decreased. Their levels then remained stable in the clinical-like stage. Second pyruvate progressively increased as PD progressed. Finally, 3 metabolites were modified only at the clinical-like stage: DMSO2 and valine decreased, while threonine increased (Figure 3D).

Thus, metabolic dysregulations observed in serum of 6-OHDA rats could be highly predictive of each PD like stage

### Brain tissue metabolic profiles in the 6-OHDA rat model reflect dysregulations in serum

Proton high resolution magic angle spinning (^1^H HRMAS) NMR spectroscopy was used to investigate the metabolic profiles of unprocessed brain biopsies (DS and Nacc) from 6-OHDA animals, sampled at the same level as the sections used for histology quantifying dopaminergic denervation. A representative spectrum of a DS biopsy from a 6-OHDA rat is shown in Figure 4A, with labeling of assigned and quantified metabolites. Regarding DS samples, the OPLS analysis (i.e. the “6-OHDA DS OPLS model”) revealed a co-evolution of cerebral metabolic profiles with the PDP score, as in serum (Figure 4B(i)). The significant dysregulations of key metabolites are illustrated in Figure 4C(i). Compared to sham animals, one metabolite, taurine, was significantly increased in the prodromal-like group, while four metabolites were significantly modified in the clinical-like group: alanine, lactate and phosphocreatine increased and glutamate decreased.

**Figure 4:**
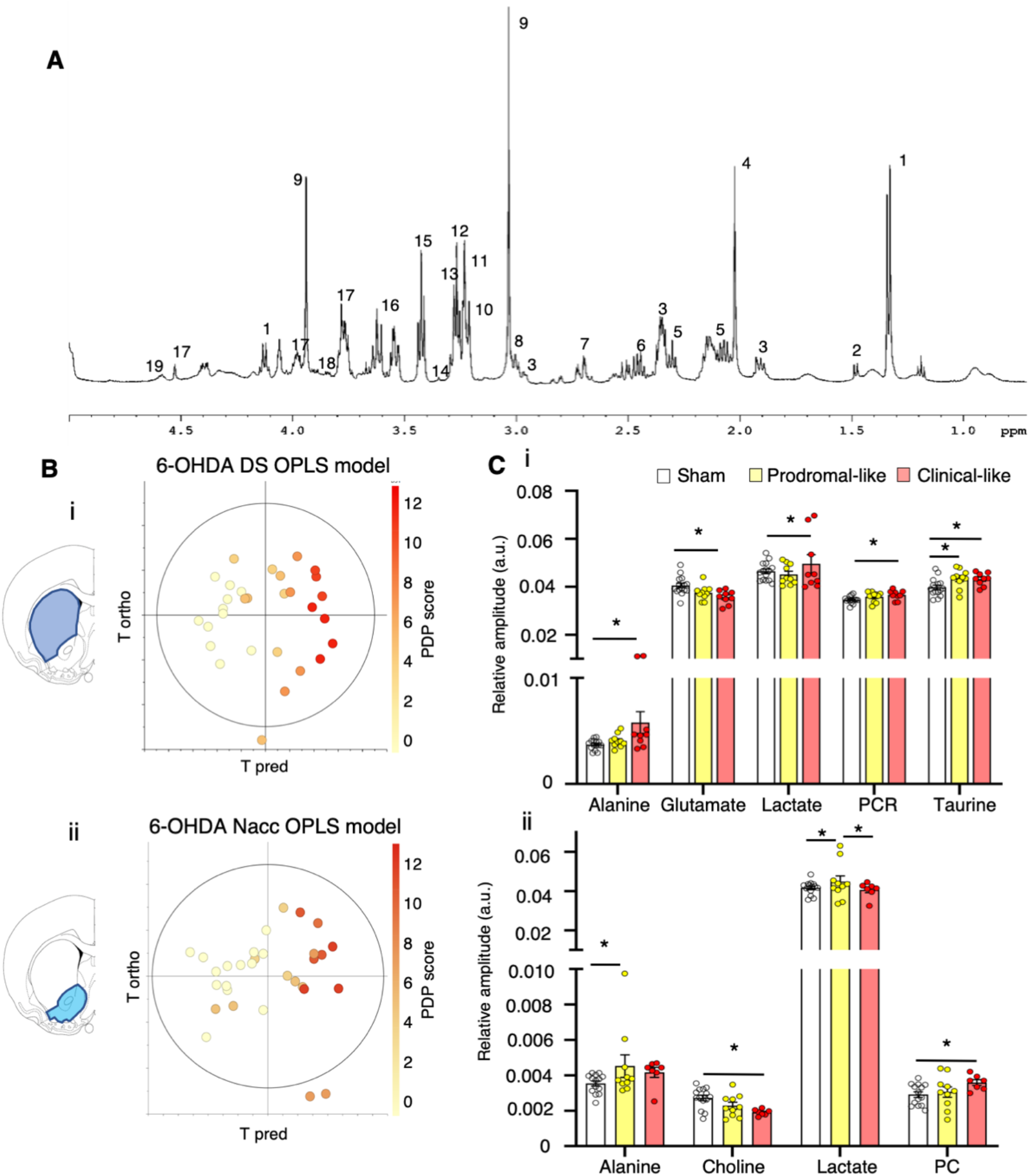
Brain metabolic profile of 6-OHDA rats is gradually modified by PD progression, consistently with serum results. (A) Example of ^1^H HRMAS NMR spectra of 6-OHDA rats brain (DS) at 500 MHz. Assignment 1-lacate; 2-alanine; 3-GABA; 4-acetate; 5-glutamate; 6-glutamine; 7-N-Acetylaspartate; 8-creatine; 9-phosphocreatine; 10-phosphoethanolamine; 11-glycerophosphocholine; 12-phosphocholine; 13-choline; 14-scyllo-inositol; 15-taurine; 16-myo-inositol; 17-ascorbate; 18-glycine; 19-glutathione. (B) Score plot of the OPLS models built with ^1^H HRMAS NMR spectra, (i) DS (the “6-OHDA DS OPLS model”) and (ii) Nacc (the “6-OHDA Nacc OPLS model”) *vs* the first predictive and the first orthogonal components. There is clear color gradation from left to right, showing that metabolic profiles evolve with PD progression, especially for DS samples (i) DS: R2Y = 0.883; Q = 0.703, 1 predictive and 3 orthogonal components, CV ANOVA p-value = 0.0003; (ii) Nacc: R2Y = 0.690; Q2 = 0.523, 1 predictive and 1 orthogonal components, CV ANOVA p-value = 0.0005. (C) Relative amplitude of metabolites in DS (i) and Nacc (ii) in sham, prodromal-like and clinical-like 6-OHDA animals for the 7 key metabolites implicated in metabolic gradation observed in the OPLS, i.e. alanine, choline, glutamate, lactate, phosphocholine (PC), phosphocreatine (PCR) and taurine. Mean ± SEM, one-way ANOVA followed by post-hoc test of Tukey and correction for multiple comparisons *: p ≤ 0.05.

The OPLS analysis of Nacc data (i.e. the “6-OHDA Nacc OPLS model”) showed the same trend, although less clear (Figure 4B(ii)). The univariate statistics revealed a significant increase of alanine and lactate in the prodromal-like group, whereas the clinical-like group was characterized by a significant increase of phosphocholine (PC) and a decrease of choline, compared to sham animals (Figure 4C(ii)).

These experiments show that both alanine and lactate exhibit similar dysregulation between serum and brain.

### Pramipexole partially reverses serum and tissue metabolic dysregulations induced by 6-OHDA

We further evaluated whether the serum metabolic profile could be influenced by the use of Pramipexole (Pra), a widely used dopaminergic treatment in newly diagnosed PD patients, and known to improve only neuropsychiatric symptoms at low doses in animal models^23, 26^. As expected, after 15 days of sub-chronic administration of Pra, the deficits in the self-administration task measured in 6-OHDA rats were fully and partially reversed in prodromal-like and clinical-like animals, respectively. In contrast, the performances of rats treated with vehicle (Veh) did not improve (Figure 5A).

**Figure 5:**
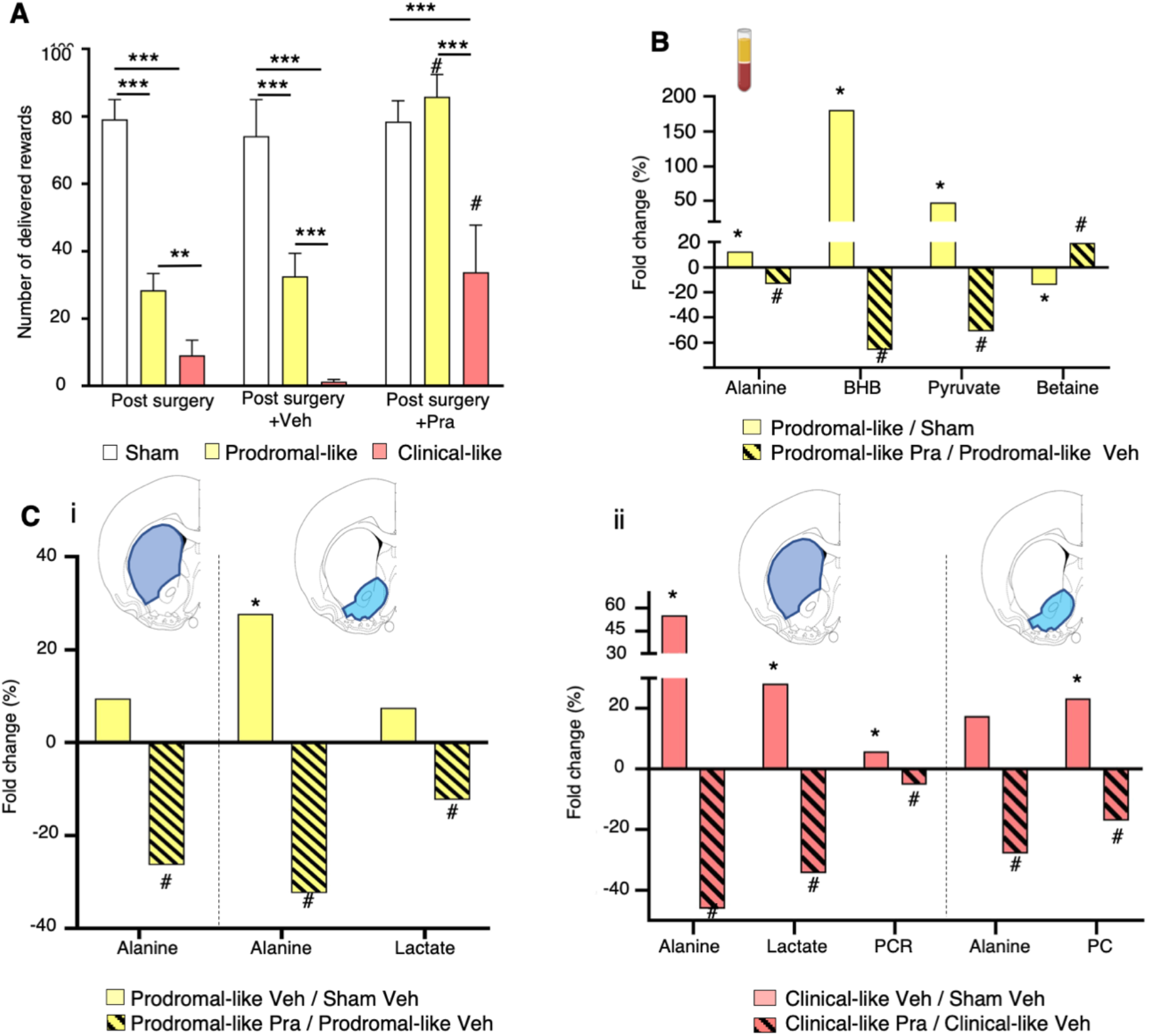
Impact of chronic treatment with a dopaminergic agonist (Pra) on behavioral and metabolic dysregulations of 6-OHDA rats. (A) *Self-administration performances*. Mean number of sucrose deliveries per session post-surgery or post chronic administration of Vehicle or Pramipexole (last 3 days). Prodromal-like animals treated with Pra show total reversion of the lesion effect on the number of sucrose deliveries compared to prodromal-like animals treated with Veh. Clinical-like animals treated with Pra show a moderate increase in the number of sucrose deliveries compared to clinical-like animals treated with Veh, without reaching the performance of sham animals. (B) *Serum metabolic dysregulations*. Percentage of variation of serum metabolites between control and prodromal-like animals (empty bar) and between prodromal-like animals treated with Veh or Pra (hatched bar). Alanine, betaine, BHB and pyruvate vary in the opposite direction compared to the lesion effect. (C) *Brain metabolic dysregulations.* Percentage of variation of brain tissue metabolites normalized to sham treated with Veh in DS and in Nacc in prodromal-like (i) and clinical-like animals (ii). We can observe the reversion of metabolic dysregulation in animals treated with pramipexole (hatched bar) compared to animals treated with Veh (full bar). Mean ± SEM, one-way ANOVA followed by post-hoc test of Tukey. *: p ≤ 0.05**: p ≤ 0.01; ***: p ≤ 0.001; #: p ≤ 0.05, prodromal-like or clinical-like Pra compared to prodromal-like or clinical-like Veh

Concerning serum metabolic profiles, to simultaneously visualize the effect of the 6-OHDA lesion and the treatment, the metabolite levels of prodromal-like animals were normalized to their levels in sham animals (lesion effect, full bar) or in lesioned animals given Veh (Pra effect, hatched bar). In prodromal-like animals, the alanine, BHB and pyruvate levels increased after lesion (Figure 3D and 5B, full bar) and were significantly decreased by Pra (Figure 5B, hatched bar), while betaine decreased after lesion, but increased in Pra animals. In the clinical-like group, metabolic reversion induced by Pra was very limited and concerned only alanine. This is consistent with the modest behavioral reversion observed (slight effect on the self-administration task and no effect on the motor task^26^) (data not shown).

In contrast to serum, for which several longitudinal samples could be taken in the same animal, brain samples could only be collected at the end of the experiment, when all animals had received chronic Pra treatment. Therefore, sham animals treated with Veh were used to normalize the metabolite levels. As in serum, Pra administration caused a decrease of some metabolites previously increased by 6-OHDA lesion. In prodromal-like animals, this was observed for alanine in DS and Nacc, and for lactate in Nacc (Figure 5C(i)). In the clinical-like group, the alanine level after Pra evolved similarly to that of prodromal-like animals, and lactate was significantly modified in DS. Furthermore, we observed a reversion of phosphocreatine and phosphocholine levels in DS and Nacc respectively (Figure 5C(ii)).

Overall, the 6-OHDA rat model shows significant alterations in both the serum and tissue metabolomes, associated with the progression of the disease and already detectable in the prodromal-like stage. Moreover, some of these alterations are partially reversed by chronic administration of Pra, which also reversed neuropsychiatric deficits.

In order to assess the specificity of the biomarkers found in the 6-OHDA model regarding progression and pathophysiological mechanisms of the disease, we extended the serum metabolomics study to two other PD animal models.

### 6-OHDA rats, alpha-synuclein rats and MPTP monkeys share common serum metabolic perturbations

First, using the same rat strain (Sprague Dawley) as in the 6-OHDA study, we selected an alpha synuclein rat model, in which the overexpression of human A53T alpha synuclein is induced in the SNc using adeno-associated viral vectors (AAV). This targeted expression leads to a progressive neurodegeneration in the targeted region, allowing the follow-up of the different stages of PD in the same animal^21^, namely a prodromal-like stage approximately 3 weeks and a clinical-like stage approximately 10 weeks after injection.

The OPLS-DA was used to evaluate whether a specific metabolic signature was associated with each stage of the disease (i.e. the “alpha synuclein serum OPLS-DA model”). Sham-animals (i.e. infected with Green Fluorescent Protein (GFP)) animals exhibited no difference between the three different time points 0, 3 and 10 weeks post injection (Figure S2), while for alpha-synuclein animals, the 3 time points were clearly separated (Figure 6A), indicating that each stage was characterized by a specific metabolic signature. In particular, an increase of BHB, glycine, pyruvate, serine and a decrease of betaine were observed over time, as in 6-OHDA rats. Moreover, a decrease of myo-inositol and an increase of acetoacetate and creatine were observed only in the alpha-synuclein model (Figure 6B).

**Figure 6:**
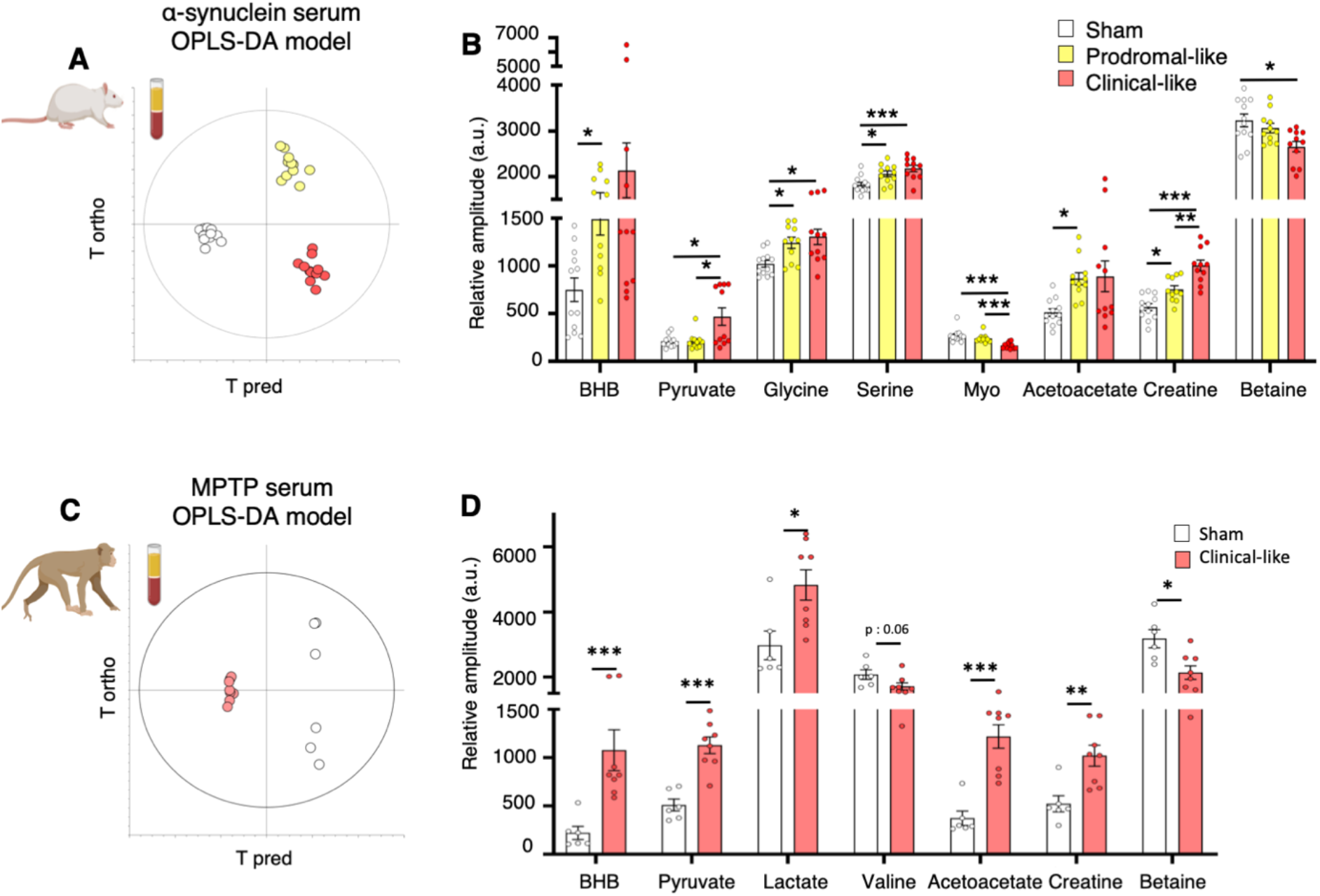
Alternative PD models from different species show similar metabolic dysregulations in serum samples. (A) Score plot of the OPLS-DA model built with ^1^HNMR spectra of serum samples from alpha synuclein rats *vs* first predictive and first orthogonal components (the “alpha-syn OPLS model”). 3 groups are clearly discriminated mimicking 3 different stages of PD (sham n = 12, prodomal-like n = 11 clinical-like n = 11). R2Y = 0.929; Q2 = 0.518; 1 predictive and 3 orthogonal components; CV ANOVA p-value = 0.008. (B) Relative amplitude of most discriminating metabolites in the “alpha-syn OPLS model”, i.e. acetoacetate, betaine, BHB, creatine, glycine, myo-inositol (myo), pyruvate, serine in sham (n = 12), prodromal-like (n = 11) and clinical-like (n = 11) animals. (C) Score plot of the OPLS-DA model built with ^1^HNMR spectra of serum samples from non-human MPTP primates *vs* first predictive and first orthogonal components (the “MPTP OPLS-DA model”). Sham (n = 6) and clinical-like (n = 8) animals are clearly discriminated. R2Y = 0.998; Q^2^ = 0.963; 1 predictive and 1 orthogonal component; CV ANOVA p-value = 0.0005. (D) Relative amplitude of most discriminating metabolites in the “MPTP OPLS-DA model” i.e. acetoacetate, alanine, betaine, BHB, creatine, lactate, pyruvate and valine in sham (n = 7) and MPTP (n = 8) animals. Mean ± SEM, mixed model followed by post-hoc test of Tukey or Mann-Whitney’s t test *: p ≤ 0.05; **: p ≤ 0.01; ***: p ≤ 0.001

These results revealed similarities in metabolic disturbances for two different PD rat models, associated with disease progression. To further investigate, we performed metabolic analysis in a non-human primate PD model, the 1-methyl-4-phenyl-1,2,3,6-tetrahydropyridine (MPTP) model, which mimics the clinical stage of PD and presents a greater homology with humans^27^. The OPLS-DA model presented in Figure 6C (i.e. the “MPTP serum OPLS-DA model”) showed clear discrimination between sham and MPTP groups. On the one hand, non-human MPTP primates presented a significant increase of lactate and a decrease of valine, like in 6-OHDA rats. On the other hand, they exhibited a significant increase of acetoacetate and creatine, like in the alpha-synuclein rats. Finally, we observed a significant increase of BHB and pyruvate, and a significant decrease of betaine, as in both rodent models (Figure 6D).

Taken together, these results reveal common metabolic alterations associated with PD progression in three different animal models, performed in two different species. To further validate the clinical relevance of these markers, we compared these preclinical results to those obtained in PD patient samples.

### Serum metabolic signatures of de novo PD patients and PD animal models show similarities

Each cohort was first analyzed individually, i.e. each *de novo* PD patient (n = 21 in each cohort) compared to an age and sex matched control (n = 30/23 NIH/Italy). Except for some metabolites, such as alanine (data not shown), the observed variations were consistent between the 2 cohorts, as shown in table 1. For instance, we observed a marginal increase of BHB and acetoacetate in Italian PD patients (+28.3%, p = 0.24 and +17.2%, p = 0.05 respectively) which was even higher in NIH PD patients (+103.2%, p = 0.09 and +28.1%, p = 0.12).

**Table 1:**
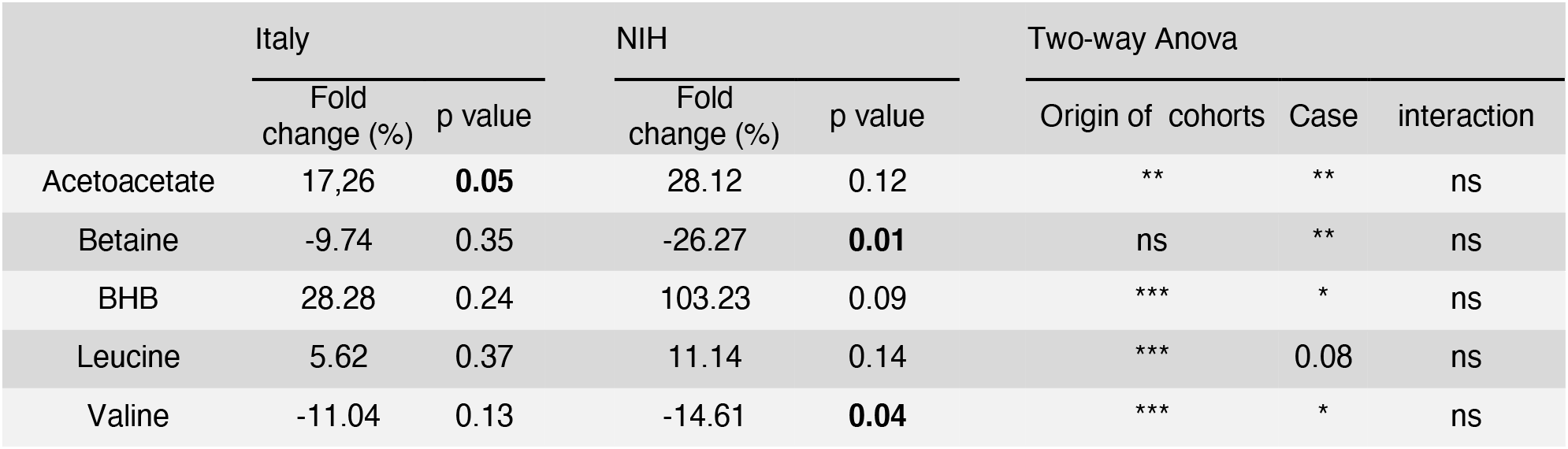
Principal individual and collective metabolite changes between PD patients and controls in Italian and NIH cohorts. Cohorts were analyzed individually by t test and together by two-way ANOVA followed by post-hoc test of Sidak *: p ≤ 0.05; **: p ≤ 0.01; ***: p ≤ 0.001; ns: non-significant

|  | Italy |  | NIH |  | Two-way Anova |  |  |
| --- | --- | --- | --- | --- | --- | --- | --- |
|  | Fold change (%) | p value | Fold change (%) | p value | Origin of cohorts | Case | interaction |
| Acetoacetate | 17,26 | **0.05** | 28.12 | 0.12 | ** | ** | ns |
| Betaine | -9.74 | 0.35 | -26.27 | **0.01** | ns | ** | ns |
| BHB | 28.28 | 0.24 | 103.23 | 0.09 | *** | * | ns |
| Leucine | 5.62 | 0.37 | 11.14 | 0.14 | *** | 0.08 | ns |
| Valine | -11.04 | 0.13 | -14.61 | **0.04** | *** | * | ns |

In order to increase statistical power, the two cohorts were pooled (PD n = 42, control n = 53). Nevertheless, as the data variance was mostly affected by sample origin, i.e Italian or NIH, independantly of the case (PD or control), we performed a two-way ANOVA with i) the origin of the cohort and ii) the case (control or PD) as factors to extract PD information only. Table 1 presents the metabolites significant for the case factor only, without interaction between the cohort origin and the case. The increase of BHB and acetoacetate observed in each cohort became significant when pooled. A significant decrease of betaine in serum of PD patients was also detected, as in the alpha synuclein and MPTP models. Moreover, a significant decrease of valine was observed as in the 6-OHDA rat model. Finally, a trend towards a decrease of leucine was noted. Moreover, the analysis of a subset of serum samples from other *de novo* PD patients treated with Pra 0.5-100 mg/day (n = 9) revealed a tendency towards normalization of BHB and betaine levels compared to non-treated PD patients, as observed in 6-OHDA rats (Figure S3). Next, we assessed the diagnostic value of the metabolites found, to discriminate prodromal-like animals from sham, or control from PD patients.

### Serum metabolites as putative composite biomarker for an early diagnosis of PD

For that, we developed multiple logistic regression models based on the metabolites significantly modified in at least three of the four groups studied (6-OHDA, Alpha synuclein, MPTP, and human), i.e. acetoacetate, betaine, creatine, BHB, pyruvate and valine. We tested each metabolite individually and all possible combinations thereof, and pooled prodromal-like animals from the 6-OHDA and alpha synuclein models.

The best regression model, as evaluated by the receiver operating characteristic (ROC) curves, retained all six metabolites, using a regression based on the following predictive algorithm (p = 1.107x10^-1^^5^):

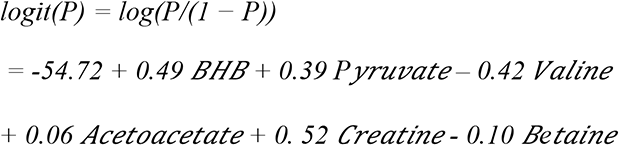

The ROC curve had an area under the curve (AUC) of 0.936. By applying the optimal threshold of 0.618 (sensitivity 0.853, specificity 0.880, Figure 7A), we found an accuracy of 92.5%, indicating that among the 40 animals included in the validation cohort (clinical-like rats and all primates), 37 were accurately predicted.

**Figure 7:**
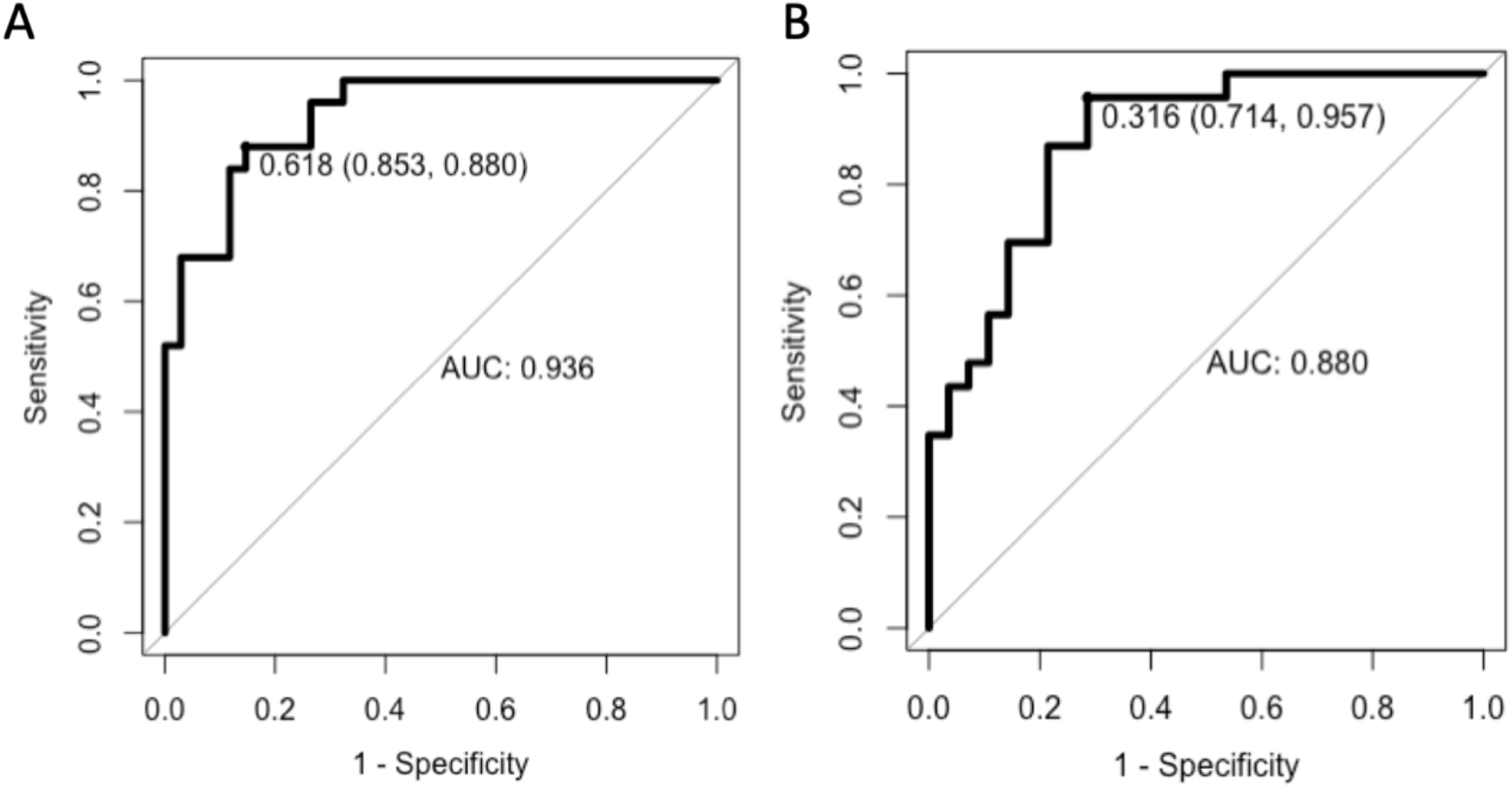
Logistic regression provided a composite biomarker built with 6 serum metabolites: BHB, acetoacetate, valine, creatine, betaine and pyruvate. (A) ROC curve with serum samples of prodromal-like and sham animal models (6-OHDA and alpha synuclein rats) (AUC = 0.936, sensitivity: 0.853, specificity: 0.88). (B) ROC curve with data of PD patients and matched controls of NIH cohort (AUC = 0.88, sensitivity: 0.957, specificity: 0.714)

Concerning human samples, we used NIH cohort as a training set since i) this is the largest cohort and ii) it includes only recently diagnosed patients (< 1 year), in contrast to the Italian cohort, which includes patients with longer PD duration (0 to 3 years post diagnosis). The latter was used as external validation set. Again, the best regression model retained all six metabolites and the following regression algorithm (p: 1.95x10^-4^):

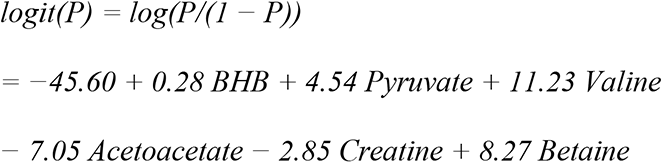

Subsequent ROC curve had an AUC of 0.88. The optimum threshold was 0.316, and corresponding sensitivity and specificity were respectively 0.957 and 0.714 (Figure 7B). The prediction of the Italian cohort resulted in 82.6% of patients being detected as PD, with 17.4% of false negatives.

Finally, a multiple logistic regression was performed using serum data from all PD animals (prodromal-like and clinical-like) from all models using the same six metabolites and was evaluated with the subsequent ROC curve presenting an AUC of 0.954 (Figure S4) and an accuracy of 88,9%.

Altogether, the accuracy of our composite biomarker from serum sample was at least 82.6% whatever the species.

## Discussion

Late and inaccurate diagnosis of PD^3, 28, 29^ takes part in too restricted patients management and therapies. The latter remain thus only symptomatic and become ineffective after several years. Although many molecules have been tested, there are still no agents or neuroprotective therapies to efficiently slow down, stop or reverse neurodegeneration in PD patients, despite promising theoretical or preclinical evidence, partly because applied too late ^30–32^. Thus, finding easily measurable and highly predictive biomarkers of the prodromal phase of the disease appears as a milestone for the acceleration of curative therapeutic development and improvement of patient care.

In the present study, using NMR-based metabolomics approaches, we observed alterations of the metabolome in serum and tissue of different animal models mimicking the different stages of PD, including the prodromal phase. NMR provides highly reproducible results and requires minimal sample preparation, which is well compatible with multicentric studies and routine in clinic. Concerning tissue samples, despite the absence of some metabolites like pyruvate and citrate, mainly due to post-mortem effects ^33^, we were able to confirm some dysregulations observed in serum, demonstrating that blood could reflect central dysfunction^34^, at least partially. In addition, the present multi-model approach allowed to overcome intrinsic limitations of each animal model and increased the incomplete predictive value of each model alone, revealing a characteristic set of dysregulated metabolites as potential biomarker. Finally, we extended the study to human serum samples from two biobanks to validate the clinical relevance of our biomarker of PD.

We have demonstrated that the profiles of 6 metabolites, acetoacetate, betaine, BHB, creatine, pyruvate, valine, combined together, could constitute an accurate PD composite biomarker, for animal models and for human samples. Using this biomarker allowed us to discriminate NIH patients from healthy controls with great sensitivity (0.95) and Italian PD patients to a slightly lesser degree (0.83). The divergent pre-analytical procedure applied for serum sampling in the two cohorts, (i.e. tubes used, sample processing…) may represent a major source of experimental variability which could explain this difference^35, 36^. Moreover, it should be noted that the human samples were collected under a very strict and standardized experimental protocol but which was not metabolomics-designed. This reflects the power of metabolomics and the robustness of our results, and supports the view that serum biomarkers, easier to use and less invasive than imaging methods, present a similar efficiency^37^.

Interestingly, among the six metabolites used in the logistic regressions to build the PD composite biomarker in animal models or in humans, the dysregulation of betaine, BHB, and pyruvate were partially corrected by chronic treatment with Pra, which also corrected the motivational deficits in prodromal-like 6-OHDA rats. This partial pharmacological reversion, also observed in *de novo* patients, demonstrates the powerful of metabolomics for identifying robust biomarkers and potentially monitoring therapeutic outcomes^38^. The metabolic dysregulations reported in the present study could provide some clues to the mechanisms impacted during PD progression (PDP). In the 6-OHDA model, the gradation of metabolic profiles was clearly associated with the evolution of the PDP score even from very early ones. In particular, glycolytic (e.g. pyruvate) and associated metabolites (e.g. lactate) were increased. Interestingly, these increases were also observed in the alpha synuclein and primate models, except for lactate in alpha-synuclein rats (Figure 8). Importantly, lactate increase was also observed in DS and Nacc and may reflect upregulation of the activity of the astrocyte-neuron lactate shuttle^39^, known to play a major role in central nervous system homeostasis and energy metabolism^40, 41^, especially in energy production. Noteworthy, alanine, intrinsically linked to this shuttle^42^, was also increased in sera and tissues of 6-OHDA animals.

**Figure 8:**
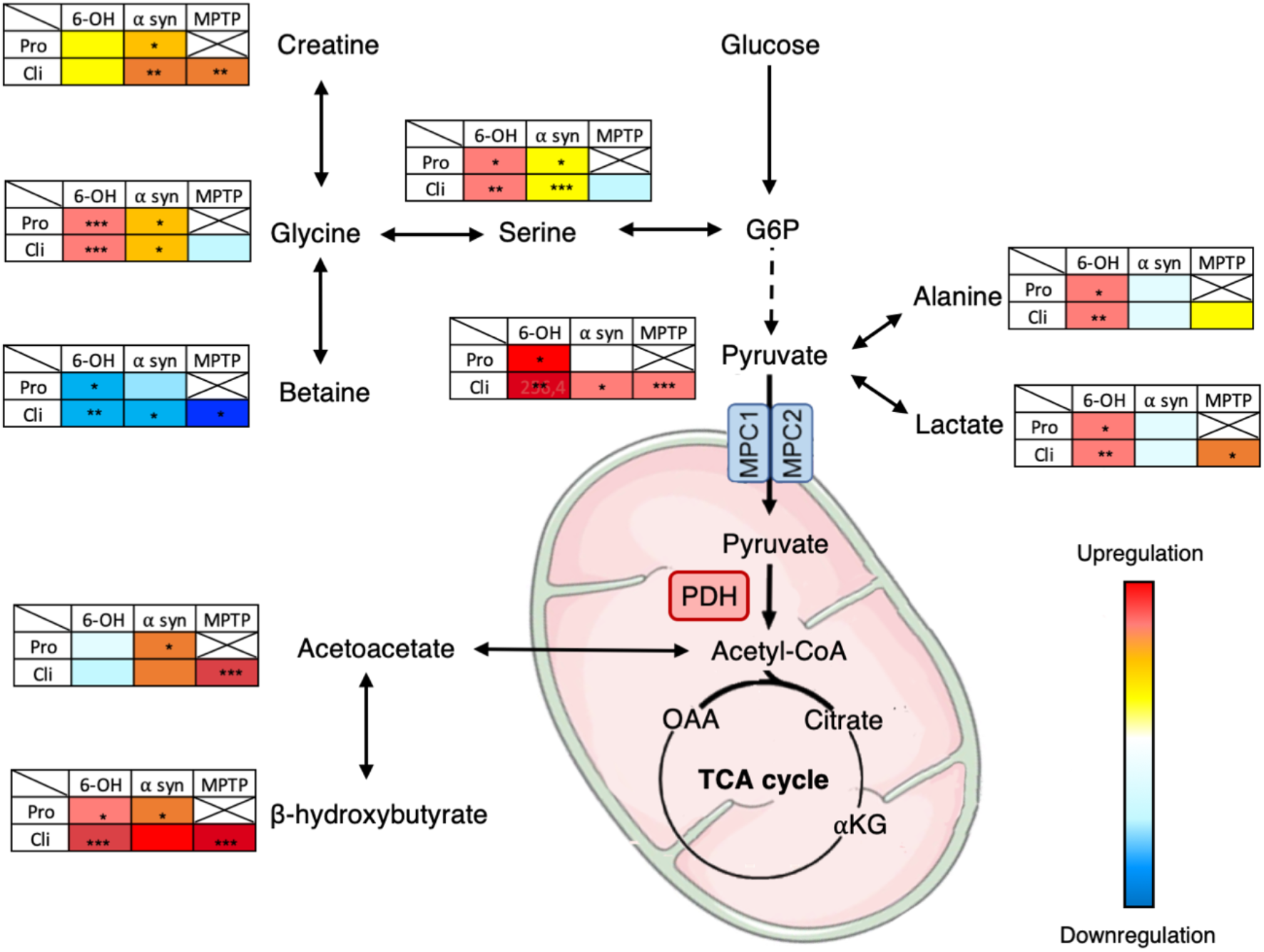
Schematic representation of altered metabolic pathways in different PD animal models mimicking different stages of the disease, suggesting a possible reprogramming of pyruvate metabolic pathway. Degrees of dysregulation were represented by color gradation compared to sham animals in each model. *: p < 0.05; **: p < 0.01; ***: p < 0.001 *Pro: Prodromal-like; Cli: Clinical-like; G6P: Glucose-6-phosphate; OAA: Oxaloacetate; αKG: alpha ketoglutarate*

Serine and glycine, possibly deriving from the first steps of glycolysis^43^, were increased in the two rat models studied. They have been described to influence mitochondrial dynamics and homeostasis^44^, and are furthermore co-agonists of NMDA receptors^45^. Their increase may support glutamatergic hyperactivity, often fundamental to neurodegeneration^46–48^. Betaine and creatine, that can derive from glycine^49, 50^, were also dysregulated but in opposite ways, i.e. betaine was decreased in all animal models, whereas creatine was increased in alpha synuclein and primate models. Both may attenuate and protect against oxidative stress^51–55^ that represents the primary cause of neuronal death in PD. In the literature, previous metabolomics studies have reported modifications of glycolysis in brain and blood samples of PD animal models^56–59^. Glycolysis has also been described as dysregulated in animal models expressing neuropsychiatric symptoms similar to those found in the early stages of PD, such as depression or anxiety^60, 61^. Together, these observations place glycolysis as a central actor^61–63^ of dysregulations occurring during PD processes, and possibly at the first stages of the disease. Surprisingly, in our study the increase of these glycolytic or glycolytic-linked metabolites (pyruvate, alanine, lactate, glycine, serine) was not associated with a rise in citrate, the first metabolite of Krebs cycle (TCA) (other metabolites of TCA cycle were not observable with the method used). These observations could suggest decoupling of glycolysis from the TCA cycle, leading to an accumulation of glycolytic metabolites. This is in agreement with the increase of ketone bodies (BHB, acetoacetate) in all animal models, which could be used as an alternative fuel to maintain TCA cycle functioning (Figure 8). In fact, ketone bodies could increase mitochondrial ATP production, and support antioxidant defenses^64^. Of note, MPTP and 6-OHDA used in animal models are known to exert their cytotoxic activities by depleting mitochondrial ATP in the brain ^65^. As such, an increase in ATP production in PD mice could protect them from developing motor symptoms^66^. Previous studies describing an increase of ketone bodies in blood of PD patients^67^ have suggested their possible neuroprotective impact^64, 68, 69^. Interestingly, high levels of ketone bodies have been associated with an inhibition of the pyruvate dehydrogenase complex (PDHC)^70^, an enzymatic complex located in the mitochondria and responsible for the transformation of pyruvate to acetyl-CoA, a crucial step between glycolysis and the TCA cycle. Such inhibition could explain the accumulation of pyruvate and associated metabolites observed in the present study^71, 72^. Moreover, down-regulation in PDHC gene expression, directly implicated in PDH activity, has been observed in the plasma of PD patients^73^. In addition, it has been shown that the mitochondrial pyruvate carrier, an integral part of the shift from glycolysis to TCA cycle by allowing the transport of pyruvate into the mitochondria, could play an important role in neuronal death and thus represent a possible target to attenuate neurodegeneration^74–77^. These two intermediaries between glycolysis and the TCA cycle may represent central players in the pathophysiological mechanisms underlying PD and need further investigation.

In summary, our study reports, for the first time, common serum metabolic alterations in three animal models mimicking PD, despite coming from different cohorts and species. Consequently, this common signature is likely to reflect specific alterations linked to PD physiopathology. In particular, we highlight specific dysregulations characteristic of the prodromal stage. Besides providing clues for deeper investigation of metabolic alterations in animal models, we also found metabolic dysregulations in sera of *de novo* PD patients (i.e. clinical phase) coming from two biobanks, not specifically designed for metabolomics. Logistic regression, using the levels of the same metabolites as those used for animals, enabled very adequate discrimination between control and PD patients in two cohorts from different origins. A more widespread, dedicated study is now necessary, particularly in prodromal and *de novo* patients.

Finally, the variations observed in the three animal models suggest possible modification in energy metabolism, especially from glycolysis to the TCA cycle. Even if this hypothesis needs additional evidence, we suggest that this alteration may be a crucial point that could be targeted to develop new PD care.

Altogether, our multi-model and translational study demonstrates the usefulness and reproducibility of untargeted metabolomics as a non-invasive approach to search for biomarkers of PD in animal models. The approach seems particularly promising for use in PD patients as serum is easily accessible and the biomarker may be relevant even at early stages of the disease when other methods fail.

## Material and methods

### Flow chart

Figure 1 illustrates a flow chart of the whole protocol applied for the animal models. The rats of the first cohort (Figure 1 (i)) were trained for 2 weeks for self-administration until they reached stable performances and were submitted to a stepping test before receiving bilateral intracerebral injection of 6-OHDA (n = 29) or NaCl (n = 22) in the SNc. After recovery and stabilization of the lesion (around 3 weeks), self-administration was resumed for one week, and the stepping test was repeated to follow the evolution of performances, before submitting each group (6-OHDA and NaCl) to 2 weeks of daily injections of Pra 0.2 mg/kg (6-OHDA Pra n = 15; NaCl Pra n = 11) or NaCl 0.09% (6-OHDA NaCl n = 14; NaCl NaCl n = 11). Self-administration was continued throughout Pra treatment and stepping test was performed for the last time at the end. Serum samples were collected after surgery, after stabilization of self-administration performances and at the end of Pra treatment. Brains were collected at the end of the behavioral procedure, snap-frozen in liquid nitrogen and kept at -80°C before being processed for histology and ^1^H HRMAS NMR experiments.

The second cohort of rats (Figure 1 (ii)), received intracerebral infusion of AAV-hA53Tα-syn (n = 12) or AAV-GFP (n = 12) in the SNc. Serum samples were taken longitudinally during the study, before and 3 and 10 weeks after AAV infusion.

Monkeys (n = 8) received intra muscular MPTP injections (0.3 to 0.5 mg/kg) every 4 to 5 days over 3 weeks^78^. Repeated low dose MPTP administration was used to mimic a moderate stage, with slow development of the disease and triggering a moderate dopaminergic lesion. Serum samples were collected at the start of the experiment and after the monkeys had reached a stable parkinsonian state (Figure 1 (iii)).

All serum samples were analyzed by ^1^H NMR at 950 MHz and brain samples were submitted to ^1^H HRMAS NMR at 500 MHz.

### Animals

#### Rats

Experiments were performed on adult male Sprague-Dawley rats (Janvier, Le Genest-Saint-Isle, France), weighing approximately 300 g (7 weeks old) at the beginning of the experiment. They were housed under standard laboratory and ethical conditions with reversed light-dark cycle (12 h/light/dark cycle, with lights ON at 7 p.m.) and with food and water available *ad libitum*.

#### Monkey

Experiments were performed on adult male *Macaca fascicularis*, weighing between 5 and 8 kg, aged between 3 and 5 years and housed under standard conditions (12 h/light/dark cycle; 23°C; 50% humidity).

Protocols complied with the European Union 2010 Animal Welfare Act and the French directive 2010/63, and were approved by the French national ethics committee (2013/113) n° 004 and by local ethical committee C2EA84 and CELYNE C2EA.

### Surgery

#### 6-OHDA bilateral injection

As previously described^20, 23^, rats were subcutaneously injected with desipramine (15 mg/kg) 30 min before 6-OHDA injection in order to protect the noradrenergic neurons. All animals were then anesthetized by intraperitoneal injection of Ketamine-Xylazine (100-7 mg/kg), and placed in a stereotactic frame (Kopf instrument). The stereotaxic coordinates of the injection sites were as follows, according to the stereotaxic atlas of Paxinos and Watson^79^ and relative to bregma: incisor bar placed at -3.2 mm: anteroposterior (AP) = -5.4 mm / lateral (L) = +/-1.8 mm / dorsoventral (V) = -8.1 mm. Animals received a bilateral injection of 2.3 µl 6-OHDA (3 mg/ml) or NaCl 0.9% (sham), at a flow rate of 0.5 µl/min. For lesioned animals presenting transient starvation 2–3 days after surgery, supplementation with a high-caloric liquid diet and palatable food was implemented for 1–2 weeks, and stopped 10 days before serum sampling. Animals that did not recover were excluded.

#### Alpha synuclein bilateral injection

As previously described^21^, rats were anesthetized with isoflurane 2.5% and placed in a stereotaxic frame (Kopf Instruments) in order to receive two bilateral injections of AAV-hA53Tα-syn (1µl-7.0×10^12^ vg/ml) or AAV-GFP (7.0 ×10^12^ vg/ml, sham animals) in the SNc (AP = -5.1 mm and -5.6 mm / L = +/-2.2 mm / V = 8 mm from bregma). After recovery from anesthesia, animals were returned to the animal facility.

### Behavioral assessment

#### 6-OHDA rats

Rats were submitted to 2 behavioral tests 3 weeks after surgery when the 6-OHDA lesion should be stabilized^80^.

##### Operant self-administration (motivational component)

Rats were trained to self-administer a 2.5% sucrose solution in operating chambers (Med Associates, St Albans, VT, USA) containing an active, reinforced lever, in which a press results in the delivery of 0.2 ml of sucrose solution associated with a light stimulus, and an inactive lever, unreinforced, for which a press causes neither delivery of sucrose nor light stimulus. This task was carried out with a fixed ratio of 1, a single support on the reinforced lever resulting in the deliverance of one reward. Each session ended when 100 rewards were obtained, or at the end of the allotted time (1 hour). The number of rewards obtained was counted for each session by MED-PC IV software.

##### Stepping test (motor component)

Animals, maintained by the posterior third of their body, were moved over a length of 90 cm by a rectilinear and regular movement from left to right and inversely along a table with a smooth surface. The number of forelimb adjustments during displacement was counted^81^. The test was carried out three times by two different experimenters, blind to the experimental conditions.

#### MPTP monkeys

The severity of parkinsonian state was evaluated using a rating scale taking into account classical motor symptoms (bradykinesia, rigidity, tremor, freezing, posture and arm posture), spontaneous activities (arm movements, spontaneous eye movements, and home cage activity) and other activities (vocalization, triggered eye movements, and feeding)^82^.

### Histology: TH-immunostaining and quantification of denervation in 6-OHDA rats

#### Brain sampling

Rats were sacrificed by decapitation 1 day after the last session of sucrose self-administration and 1 hour after the last Pra or placebo injection (i.e., 6 weeks after 6-OHDA infusion or saline). Brains were immediately frozen in liquid nitrogen and stored at -80 °C. They were then processed at -20 °C for both immunostaining and HRMAS NMR to further match metabolomics and immunostaining-based lesions. For HRMAS NMR, thick sections of DS and Nacc tissue were pooled into disposable inserts, and kept at -80°C until NMR analysis. For immunostaining, 14 µm coronal sections of striatal levels of interest^20^ were sampled and stored at -20 °C. Immunostaining was then carried out as previously described^20, 23^. Briefly, after post-fixation with paraformaldehyde 4%, striatal slices were incubated with an anti-TH antibody (mouse monoclonal MAB5280, Millipore, France, 1: 2500) overnight at 4°C. Then slices were incubated with biotinylated goat anti-mouse IgG antibody (BA-9200, Vector Laboratories, Burlingame, CA, USA; 1: 500). Avidin-peroxidase conjugate revealed immunoreactivity (Vectastain ABC Elite, Vector Laboratories Burlingame, CA, USA).

Quantification of the extent of striatal dopaminergic denervation was determined with ICS FrameWork computerized image analysis system (Calopix, 2.9.2 version, TRIBVN, Châtillon, France) coupled with a light microscope (Nikon, Eclipse 80i). After drawing masks from three striatal levels, optical densities (OD) were measured for each striatal sub-region (DS and Nacc). OD were expressed as percentages relative to the mean optical density obtained from the homologous regions of sham-operated animals.

### PD progression score in 6-OHDA model

In order to assess PD Progression (PDP) in 6-OHDA rats, we developed a scale, referred to as the “PDP score”, which combines 3 criteria: 1) evaluation of neuropsychiatric symptoms (i.e. apathetic like behavior), 2) evaluation of motor symptoms (i.e. fine motor deficit), both of them used in clinic, and 3) level of dopaminergic denervation in DS, only available *post-mortem* in the patient. Each criterion was quantified for each animal, with values ranging from 0 to 4, as illustrated in Figure 2a. They were summed to produce the PDP score, ranging from 0 to 12. For the evaluation of behavioral deficits in the self-administration and stepping tests (criteria 1 and 2, respectively), we compared the performances of each animal before and after surgery. A value of 0 for each of these two criteria corresponds to an absence of deficit, or to a deficit lower than 30%, i.e. the normal daily variability in the tests used. The following subcategories were set as: 1 = small deficit (30-50%); 2 = medium symptoms (50-70%); 3 = strong symptoms (70-90%); 4 = large or total deficit (> 90%).

Considering the fact that motor symptoms appear after 70% of striatal dopaminergic neurons loss in PD patients, the value for dopaminergic denervation in the DS (criterion 3) was used as the limit between the prodromal and the clinical phases.

Regarding the PDP score obtained by addition of the values of the 4 criteria, 6-OHDA animal was classified as: 1) asymptomatic rats with weak lesions but no behavioral symptom (PDP score = 1 to 2), 2) prodromal-like animals i.e. presenting only neuropsychiatric disorders and a small DS lesion (PDP score = 3 to 7) and 3), clinical-like animals i.e. presenting neuropsychiatric disorders, motor symptoms and widespread DS lesion (PDP score = 8 to 12). Shams were attributed a 0 score.

Considering the low number of asymptomatic animals and the fact that they do not present any clinical relevance, they were excluded from the study.

### Human cohorts

Blood samples were obtained from PD patients and matched-control subjects from the Parkinson’s Disease Biomarkers Program (PDBP) Consortium, supported by the National Institute of Neurological Disorders and Stroke at the National Institutes of Health (NIH-USA) and from Santa Lucia Fondazione (Rome, Italy) cohorts. All donors gave written informed consent.

Inclusion criteria for NIH patients included recent diagnosis (< 1 year) of PD according to the new criteria of the Movement Disorders Society (MDS). For the Italian cohort, diagnosis criteria were less restrictive and patients diagnosed for longer times than the NIH cohort were included (0 to 3 years post diagnosis). In the two cohorts, patients had not received any anti-parkinsonian treatment at time of inclusion, presented no dementia or active psychiatric disorders nor any other medical condition that could compromise the study. Inclusion criteria for controls were absence of neurological diseases, no family history of movement disorders, and no specific medical conditions.

The cohort from NIH included 21 PD patients without antiparkinsonian treatment, 9 PD patients who received Pra treatment and 30 age and sex-matched controls. The Santa Lucia cohort comprised 21 PD patients and 23 age and sex-matched controls. All clinical and demographic information collected are summarized in supplemental Table 1.

### NMR experiments

#### ^1^H HRMAS NMR in 6-OHDA rat brain samples

Just before HRMAS analysis, 10 µl D_2_O was added to the inserts containing the brain biopsies, which was then sealed and packed into a 4 mm zirconia MAS rotor.

#### Data acquisition

All HRMAS NMR spectra were acquired, as previously described^83^, on a Bruker (Billerica, USA) Advance III spectrometer (IRMaGe, CEA Grenoble, France) at 500 MHz. Samples were spun at 4000 Hz at the magic angle (54.7°) and temperature maintained at 4°C for all experiments. One-dimension spectra were acquired using a Carr-Purcell-Meiboom-Gill (CPMG) pulse sequence (TE = 30 ms, 256 averages, 17 min). The residual water signal was pre-saturated during 1.7 s of relaxation.

#### Data processing

Quantification was performed with jMRUI software using quantitation based on a quantum estimation (QUEST) procedure^84^. This procedure requires the use of a metabolite database and a complete assignment of spectra. Nineteen metabolites were assigned and quantified: acetate, alanine, ascorbate, choline, creatine, γ-aminobutyrate (GABA), glutamate, glutamine, glycine, glycerophosphocholine, glutathion, lactate, myo-inositol, *N*-acetylaspartate, phosphocholine, phosphocreatine, phosphoethanolamine, scyllo-inositol, and taurine. The amplitude of metabolite calculated by QUEST was normalized to the total spectrum signal. CRLB were calculated for each metabolite as estimates of the standard deviation of the fit.

#### ^1^H NMR of serum

##### Serum sampling

For rats, blood was collected under gas anesthesia with isoflurane (2%) from the caudal vein after 2 hours of fasting, and was stored in ice before being rapidly centrifuged at 1600g for 15 min at 4° C. The supernatant serum was removed and stored at -80°C until the day of NMR. The time before freezing never exceeded 30 min^85^.

For monkeys, blood was collected under anaesthesia (atropine 0.05 mg/kg i.m. followed by zoletil 15 mg/kg i.m.) from the saphenous vein after a 12-hour fast and submitted to the same protocol as rat blood.

For humans, blood was collected after an overnight fast, stored at room temperature for 15 to 60 min before being centrifuged between 1200 and 1500 g at 4°C during 10 to 15 min. The supernatant serum was then removed and stored at -80 °C.

The day of NMR experiments, samples were slowly thawed on ice, then quickly centrifuged to eliminate possible cryoprecipitates. NMR tubes were filled with 60 µl serum sample and 120 µl phosphate buffer saline (PBS) 0.1 M in D_2_O (50% of D_2_O, pH = 7.4), and stored at 4 °C until NMR acquisition.

##### Data acquisition

All animal and human serum samples were submitted to the same NMR protocol. ^1^H NMR experiments were performed on a Bruker Advance III NMR spectrometer at 950 MHz (IBS, Grenoble, France) using a cryo-probe with a 3 mm tube holder. One-dimension spectra were systematically recorded using a CPMG pulse sequence for edition of metabolites. The residual water signal was pre-saturated during 2 seconds of relaxation.

Assignment of peaks was performed using 2-dimension homonuclear ^1^H-^1^H (TOCSY) and heteronuclear ^1^H-^13^C (HSQC) spectra of selected samples, and database. When necessary, addition of selected metabolites was used to unravel ambiguous assignments.

##### Data processing

The free induction decays were Fourier transformed and manually phased with the Bruker software Topspin version 3.6.2. Then, further pre-processing steps (baseline correction, alignment, bucketing) were performed using NMRProcFlow v1.2 online (http://nmrprocflow.org).The spectra were segmented in 0.001 ppm buckets between 0 and 10 ppm, with exclusion of residual water peaks, macromolecule signals and other regions corresponding to pollution (δ5.0–4.7; δ5.25–5.38; δ2.2–2.0; δ1.4–1.25; 1.15-1; δ0.9–0.5). Each bucket was normalized to the total sum of buckets.

### Statistical analysis

#### Multivariate analysis

Data from liquid or HRMAS NMR were imported into SIMCA v.14 (Malmö, Sweden) for multivariate statistics. An unsupervised Principal Components Analysis (PCA) was first used for global visualization of the distribution of all samples, followed by an OPLS to find discriminatory metabolites associated with a specific stage of the disease. For this latter, either a continuous variable (i.e. PDP score), or a discrete variable (i.e. group) was used to label each sample, leading to OPLS-DA (discriminant analysis) in this later case. The total number of components was determined using the cross-validation procedure, which produces the R2Y and Q2 factors that indicate respectively the goodness of the fit and the predictability of the model. A model is considered as robust and predictive when both factors are ≥0.5. Scores were plotted in 2D *vs* the two first components of the OPLS models while loadings were plotted in 1D to mimic an NMR spectrum but with positive/negative peaks indicating respectively up/down regulated metabolites. Furthermore, in this “statistical spectrum”, each NMR variable was color-coded according to their correlation with group belonging, with a “hot” colored (*e.g.*, red) metabolite being more significant than a “cold” colored (*e.g.*, green) one. Metabolites with correlation ≥0.5 were considered the most discriminant and were submitted to further univariate analysis. Analysis of variance of cross-validated predictive residuals (CV-ANOVA) was used to assess the significance of the model.

The metabolites showing significant modification in at least three groups among the four studied (6-OHDA, Alpha synuclein, MPTP, and human) were submitted to multiple logistic regression in order to evaluate their relevance as a potential predictive biomarker of PD using R software (version 3.6.1 – R core team). Logistic regression were presented using generic equation of the form: *log(P/(1 − P)) = β_0_ + β_1_ * x*, where x represents metabolite relative amplitude and β_0_ and β_1_ are the parameters associated with the intercept and the degree of change in metabolite, respectively. The regressions, including cross validation, were conducted for prodromal-like animals (6-OHDA and alpha synuclein prodromal-like rats) and clinical-like animals were used for external validation (6-OHDA and alpha synuclein clinical-like rats, and all primates). The same analyses were conducted for the NIH cohort. The Italian cohort was used for external validation. All combinations of the retained metabolites were tested, and subsequent ROC curves were generated for each of them to assess the quality of the fit using AUC and optimal threshold which maximizes sensitivity and specificity.

#### Univariate analysis

All univariate analyses were performed using Graphpad Prism 8 software (San diego–USA). All results were expressed as mean values ± standard error of mean (SEM) with a threshold for significance fixed at 0.05.

##### 6-OHDA models

For self-administration and stepping tests, Pra effect univariate analysis were performed by applying repeated measure (RM) one-way ANOVA followed by a Sidak post hoc test.

For histological and metabolomics data, one-way ANOVA followed by post hoc Tukey test with correction for multiple comparisons were performed.

##### Alpha synuclein models

For longitudinal alpha synuclein study, some values were missing due to artefacts during NMR measurement. Data were therefore analyzed by fitting a mixed model proposed using a compound symmetry covariance matrix, and were fitted using Restricted Maximum Likelihood (REML). This model was followed by a post hoc Tukey test.

##### MPTP monkeys

Considering the low number of animals, the non-parametric Mann–Whitney test was used.

##### Humans

*Data* were log-transformed given the non-Gaussian distribution of metabolite levels. First, t-tests were performed individually in the two different cohorts. Then we applied two-way ANOVA with i) origin of the cohort and ii) group (control or PD) as factors followed by the post hoc Sidak test.

## Acknowledgements

Data and biospecimens used in the present study were obtained from the Parkinson’s Disease Biomarkers Program (PDBP) Consortium, supported by the National Institute of Neurological Disorders and Stroke at the National Institutes of Health. Investigators include: Roger Albin, Roy Alcalay, Alberto Ascherio, Thomas Beach, Sarah Berman, Bradley Boeve, F. DuBois Bowman, Shu Chen, Alice Chen-Plotkin, William Dauer, Ted Dawson, Paula Desplats, Richard Dewey, Ray Dorsey, Jori Fleisher, Kirk Frey, Douglas Galasko, James Galvin, Dwight German, Lawrence Honig, Xuemei Huang, David Irwin, Kejal Kantarci, Anumantha Kanthasamy, Daniel Kaufer, James Leverenz, Carol Lippa, Irene Litvan, Oscar Lopez, Jian Ma, Lara Mangravite, Karen Marder, Laurie Orzelius, Vladislav Petyuk, Judith Potashkin, Liana Rosenthal, Rachel Saunders-Pullman, Clemens Scherzer, Michael Schwarzschild, Tanya Simuni, Andrew Singleton, David Standaert, Debby Tsuang, David Vaillancourt, David Walt, Andrew West, Cyrus Zabetian, Jing Zhang, and Wenquan Zou. The PDBP Investigators have not participated in reviewing the data analysis or content of the manuscript

The authors would like to thank Adrien Favier and Alicia Vallet from IBS (Grenoble) for technical assistance concerning NMR experiments and IR-RMN-THC Fr3050 CNRS for financial support for conducting the research. They also thank Pierre Alain Bayle from CEA (Grenoble), for his help concerning HRMAS experiments. Finally, we thank Jacques Brocard for his help and advices concerning univariate statistics and logistic regression methods as well as Fiona Hemming, Yvan Vachez and Sara Meoni for English corrections and critical reading of the manuscript.

## Author contributions

Research: D.M, T.D, C.C, S.C, E.L.B, S.B, F.F Conducted experiments: DM, TD, CC, MD, MBM Provide samples: P.O.F, V.S, PB Designed research: D.M, S.B, F.F Analysed data D.M, S.B, F.F Wrote the manuscript D.M, S.B, F.F

## Abbreviations

6-OHDA: 6-hydroxydopamine
AUC: Area under the curve
DS: Dorsal striatum
GFP: Green fluorescent protein
HRMAS: High resolution magic angle spinning
MPTP: 1-methyl-4-phenyl-1,2,3,6-tetrahydropyridine
Nacc: Nucleus accumbens
NMR: Nuclear magnetic resonance
OPLS: Orthogonal partial least square
OPLS-DA: Orthogonal partial least square discriminant analysis
PD: Parkinson’s disease
PDH: Pyruvate dehydrogenase
PDHC: Pyruvate dehydrogenase complex
PDP: Parkinson’s disease progression
Pra: Pramipexole
ROC: Receiver operating characteristic
SNc: Substantia nigra pars compacta
TH-IR: Tyrosine hydroxylase immunoreactivity
Veh: Vehicle

