## supplemental material for "Early diagnosis of Parkinson’s disease: A cross-species biomarker"

### Supplemental data

Supplemental table 1

| | NIH | | Italy | |
|---|---|---|---|---|
| | Control | PD | Control | PD |
| Age (years) | 61.4 ± 1.8 | 62,0 ± 2.1 | 57.26 ± 1.89 | 60.38 ± 2.00 |
| Sex | 16 M / 14 W | 9 M / 12 W | 9 M / 14 W | 14 M / 7 W* |

**Supplemental table 1: Demographic characteristics of two Human cohorts**.
* 2 patients sex unknown
Data are presented by mean value ± SEM. *M: Men ; W: Women*

### Supplemental Figure 1

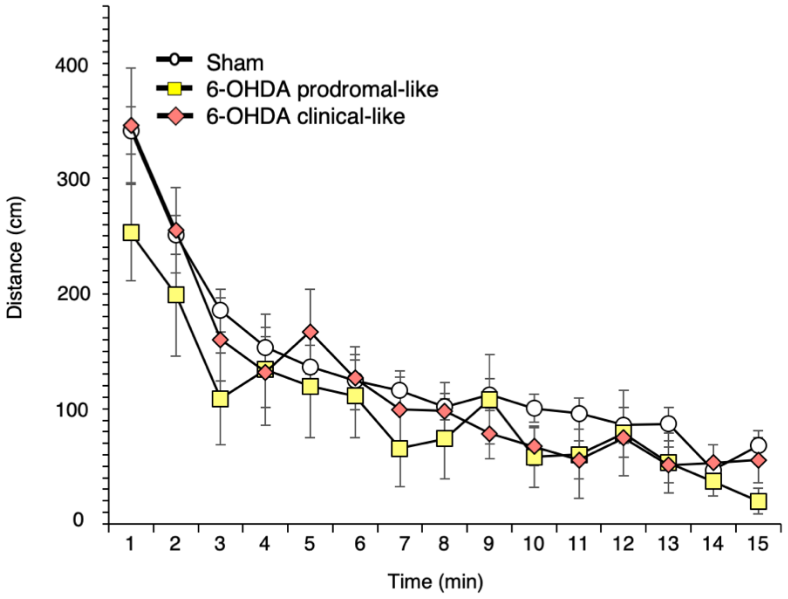


**Supplemental figure 1: Ambulatory motor activity measured by open field.**
Dopaminergic lesions did not affect horizontal ambulatory activity during a 15-min period in an open area.
Data are presented by mean value of each minute ± SEM.

Supplemental table 2

| Metabolite | Group | $^{1}$H ppm | $^{13}$C ppm | Multiplicity |
|---|---|---|---|---|
| 3-hydroxybutyrate | ɣ-$CH_3$ | 1.19 | 24,4 | d |
| | half α-$CH_2$ | 2.30 | 49,2 | dd |
| | half α-$CH_2$ | 2.39 | 49,2 | dd |
| | β-CH | 4.15 | | m |
| Acetoacetate | | 3,44 | | s |
| | | 2.27 | | s |
| Acetate | $CH_3$ | 1.91 | 25.9 | s |
| Acetone | $CH_2CO$ | 2.22 | 32,9 | s |
| Alanine | $CH_3$ | 1.47 | 18.8 | d |
| | α-CH | 3.77 | 53.5 | q |
| Albumin lysyl | ϵ-$CH_2$ | 2.88 | 42.0 | t |
| | ϵ-$CH_2$ | 2.95 | 41.9 | t |
| | ϵ-$CH_2$ | 3.01 | 42.0 | t |
| Arginine | ɣ-$CH_2$ | 1.64 | | |
| | ɣ-$CH_2$ | 1.69 | | |
| | β-$CH_2$ | 1.89 | | |
| | δ-$CH_2$ | 3.23 | 43.2 | t |
| Aspartate | half β-$CH_2$ | 2.66 | | q |
| | half β-$CH_2$ | 2.80 | | dd |
| Betaine | CH2 | 3,9 | | s |
| | $CH_3$ | 3.26 | | s |
| Cholesterol | C18 (in HDL) | 0.66 | | m |
| | C18 (in VLDL) | 0.68 | | |
| | C26 and C27 | 0.83 | 25.2 | m |
| Choline | N $(CH_3)_3$ | 3.21 | 56.7 | s |
| | | 3.51 | | |
| | | 4.06 | | |
| Citrate | half $CH_2$ | 2.53 | | d |
| | half $CH_2$ | 2.66 | | d |
| Creatine | $CH_3$ | 3.03 | | s |
| | $CH_2$ | 3.92 | | s |
| Creatinine | $CH_3$ | 3.04 | | s |
| | $CH_2$ | 4.05 | | s |
| Dimethylamine | $CH_3$ | 2.71 | | s |
| Ethanol | $CH_3$ | 1.17 | 21.6 | t |
| | $CH_3COH$ | 3.65 | | q |
| Fatty acids (mainly LDL) | $CH_3(CH_2)n$ | 0.84 | 16.5 | m |
| | $(CH_2)n$ | 1.27 | 32.2 | m |
| Fatty acids (mainly VLDL) | $CH_3CH_2CH_2C=$ | 0.86 | | m |
| | CH2CH2CO | 1.57 | 27.4 | m |
| | $CH_2CH_2CH_2CO$ | 1.29 | | m |
| Fatty acids | $CH_3CH_2$ | 0.93 | 21.05 | m |
| | $CH_3CH_2(CH_2)n$ | 1.24 | 34.4 | m |
| | $CH_3CH_2(CH_2)n$ | 1.26 | 25.2 | m |
| | $CH_2$ | 1.26 | 19.2 | m |
| | $CH_2$ | 1.30 | | m |
| | $CH_2CH_2C=C$ | 1.68 | 29.2 | |

| Metabolite | Group | $^{1}H$ ppm | $^{13}C$ ppm | Multiplicity |
|---|---|---|---|---|
| Fatty acids | $CH_2C=C$ | 2.00 | 29.7 | m |
| | $CH_2CO$ | 2.22 | 36.3 | m |
| | $C=CCH_2C=C$ | 2.72 | 28.1 | m |
| | CH=CHCH2CH=CH | 5.26 | 130.6 | m |
| | CH=CHCH2CH=CH | 5.29 | 132.2 | m |
| Formate | CH | 8.45 | | s |
| Fructose | | 3,99 | | m |
| | | 4.01 | | dd |
| Fucose / β-Galactose | | 4.54 | | d |
| Glucose | H2 | 3.24 | 76.9 | t |
| | H4 | 3.40 | 72.4 | t |
| | H4 | 3.41 | 72.4 | t |
| | H5 | 3.46 | 78.6 | m |
| | H3 | 3.48 | 78.5 | t |
| | H2 | 3.53 | 74.3 | q |
| | H3 | 3.71 | 75.6 | t |
| | half $CH_2$-C6 | 3.72 | 63.5 | q |
| | half $CH_2$-C6 | 3.76 | 63.4 | m |
| | H5 | 3.82 | 74.2 | ddd |
| | half $CH_2$-C6 | 3.84 | 63.4 | m |
| | half $CH_2$-C6 | 3.89 | 63.5 | dd |
| | H1 | 4.64 | 98.7 | d |
| | H1 | 5.23 | 94.9 | d |
| Glutamate | half β-$CH_2$ | 2.04 | | m |
| | half β-$CH_2$ | 2.12 | | m |
| | half γ-$CH_2$ | 2.34 | 33.9 | m |
| | half γ-$CH_2$ | 2.36 | | m |
| | | 3.74 | | m |
| Glutamine | | 2.08 | | |
| | | 2.09 | | |
| | half β-$CH_2$ | 2.11 | 29.7 | m |
| | half γ-$CH_2$ | 2.44 | 33.9 | m |
| | | 2.46 | 57.4 | m |
| | | 3.74 | | |
| Glycerol | half $CH_2$ | 3.56 | 65.8 | q |
| | half $CH_2$ | 3.65 | 65.6 | q |
| | $C_2$-H | 3.87 | 74.6 | m |
| Glycerophosphocholine | | 3.22 | | s |
| | $NCH_2$ | 3.66 | 68.7 | m |
| | $OCH_2$ | 4.29 | 62.2 | m |
| Glycerol backbone | | 4.06 | | |

| Metabolite | Group | $^{1}H$ ppm | $^{13}C$ ppm | Multiplicity |
|---|---|---|---|---|
| PGLYs and TAGs | CHOCOR | 4.22 | | |
| | | 5.20 | | |
| Glycine | $CH_2$ | 3.55 | 44.3 | s |
| Histidine | | 3.09 | | dd |
| | | 3.98 | | dd |
| | H4 | 7.04 | | s |
| | H2 | 7.75 | | s |
| Isoleucine | δ-$CH_3$ | 0.93 | 13,9 | t |
| | β-CH3 | 1.00 | 17,15 | d |
| | half γ-$CH_2$ | 1.24 | | |
| | half γ-$CH_2$ | 1.46 | | |
| | | 1.96 | | |
| | | 3.65 | | |
| Lactate | $CH_3$ | 1.32 | 22.7 | d |
| | CH | 4.11 | 71.2 | q |
| Lactose | | 3.55 | | |
| | | 3.66 | | |
| | | 3.97 | | |
| | | 4.45 | | |
| Leucine | δ-$CH_3$ | 0.95 | | d |
| | δ-$CH_3$ | 0.96 | | d |
| | | 1.66 | | m |
| | | 1.70 | 42.7 | m |
| | | 1.73 | | m |
| | α-CH | 3.71 | | |
| Lysine | γ-$CH_2$ | 1.43 | | m |
| | γ-$CH_2$ | 1.49 | | m |
| | δ-$CH_2$ | 1.72 | | m |
| | β-$CH_2$ | 1.88 | | m |
| | β-$CH_2$ | 1.91 | | m |
| | | 3.02 | | t |
| | | 3.74 | | t |
| Mannose | | 4.89 | | d |
| | | 5.18 | | d |
| Methanol | $CH_3OH$ | 3.35 | | s |
| Methionine | | 2,64 | 31,4 | t |
| Methionine | | 2.15 | | s |
| Myo-inositol | | 4.05 | | |
| | | 3.27 | | t |
| N-acetyl-glycoprotein 1 | $NHCOCH_3$ | 2.04 | 24.7 | s |
| N-acetyl-glycoprotein 2 | $NHCOCH_3$ | 2.07 | 25 | s |

| Metabolite | Group | $^{1}$H ppm | $^{13}$C ppm | Multiplicity |
|---|---|---|---|---|
| Phenylalanine | half β-$CH_2$ | 3.26 | | |
| | α-CH | 3.97 | | |
| | H2, H6 | 7.31 | | d |
| | H4 | 7.35 | | m |
| | H3, H5 | 7.40 | | m |
| Proline | γ-$CH_2$ | 1.98 | | m |
| | γ-$CH_2$ | 2.01 | | m |
| | half β-$CH_2$ | 2.05 | | m |
| | half β-$CH_2$ | 2.34 | | m |
| | half δ-$CH_2$ | 3.33 | | m |
| | α-CH | 4.12 | | m |
| Succinate | | 2.39 | | s |
| Threonine | γ-$CH_3$ | 1.31 | | d |
| | α-CH | 3.55 | | d |
| | β-CH | 4.23 | | m |
| Trehalose | | 3.40 | 74.3 | |
| Tyrosine | | 6.88 | | d |
| | H2, H6 | 7.18 | | d |
| Valine | CH3 | 0.98 | 19.4 | d |
| | CH3 | 1.03 | 20.6 | d |
| | β-CH | 2.26 | | m |
| | α-CH | 3.60 | 63.4 | d |
| Urea | $NH_2C{=}ONH_2$ | 5.77 | | |
| Xylose | | 3.41 | 78.6 | |

**Supplemental table 2: List of metabolites identified in serum samples.**

For each metabolite, chemical group, assignment for $^{1}$H and $^{13}$C and multiplicity of peak are presented.

Multiplicity: singulet (s), doublet (d), doublet doublet (dd), multiplet (m)

#### Supplemental Figure 2

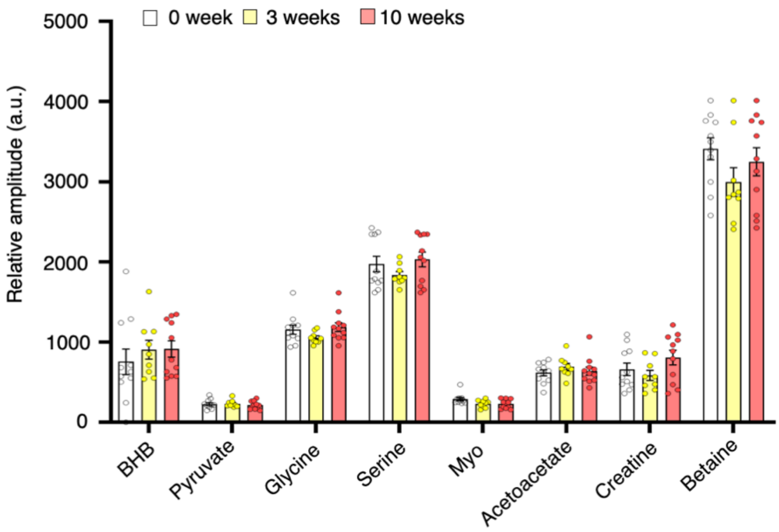


**Supplemental figure 2: GFP viral infusion does induce metabolic dysregulation in serum samples.**

Representative histogram showing relative quantitative variations of signal between different samples in GFP animals for acetoacetate, betaine, BHB, creatine, glycine, myo-inositol, pyruvate, serine which represent key metabolites implicated in discrimination of 3 groups in OPLS-DA of alpha syn animals. White bar corresponds to samples at week 0, yellow to 3 weeks post GFP infusion and red to 10 weeks after the same infusion. Data are presented as mean value ± SEM and tested by one-way ANOVA followed by posthoc test of Tuckey.

#### Supplemental Figure 3

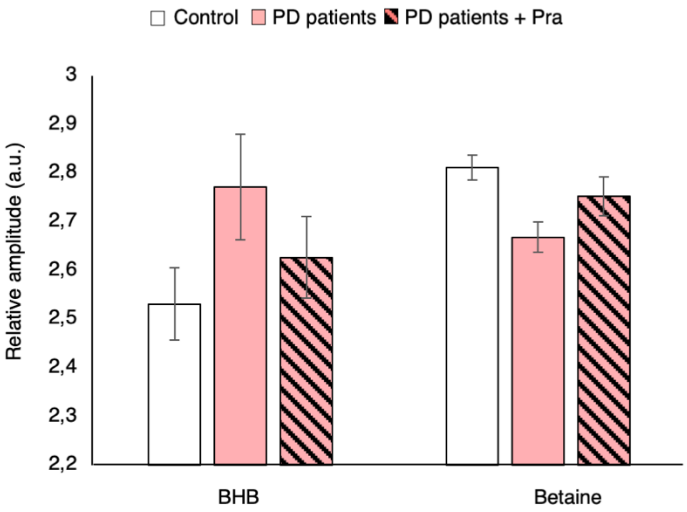


**Supplemental figure 3: BHB and betaine levels appear partially normalized in NIH PD patients treated with pramipexole.**

Representative histogram showing relative quantitative variations of signal between Control (white bar), PD patients (red bar) and PD patients treated with Pra (hatched bar) for BHB and betaine. Data are presented as mean value ± SEM.

#### Supplemental Figure 4

A

$$\mathrm{logit}(P) = \log(P/(1-P))$$

$$= 132.29 - 0.45\,BHB - 0.82\,Pyruvate + 0.36\,Valine + 0.17\,Acetoacetate - 0.39\,Creatine + 0.07\,Betaine$$

B

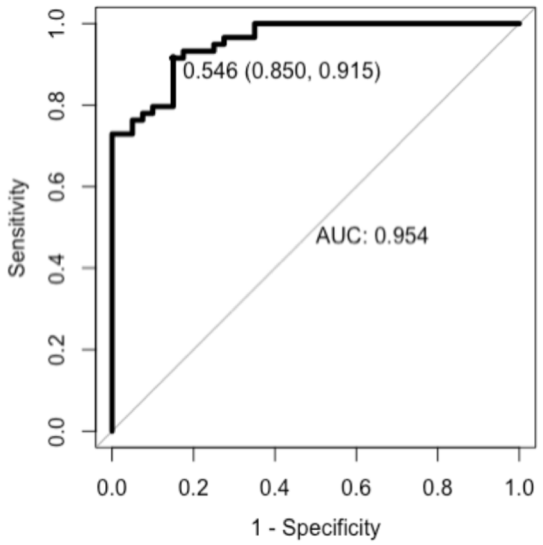


**Supplemental figure 4: Logistic regression curve for panel of serum metabolites included : BHB, acetoacetate, valine, creatine, betaine and pyruvate.**

(A) ROC curve from serum samples of all PD-like animal models. AUC = 0.954, sensibility : 0.85, specificity : 0.915.

(B) Algorithm of logistic regression for all PD-like animals vs sham.
